# Baseline expression of c-Myc defines the tissue specificity of oncogenic K-Ras

**DOI:** 10.1101/2025.07.21.665965

**Authors:** Olesja Popow, Qing Yu, Clarance Yapp, Shannon Hull, João A Paulo, Benjamin Hanna, Shidong Xu, Yanan Kuang, Carlie Sigel, Cloud P Paweletz, Steven P Gygi, Kevin M Haigis

## Abstract

*KRAS* is among the most frequently mutated oncogenes in cancer. Yet, mutations in *KRAS* are common only in tumors originating from a subset of tissues. It is critical to understand the molecular mechanisms underlying this oncogene tissue specificity. Utilizing genetically engineered mouse models carrying a conditional oncogenic allele of *Kras*, we expressed activated K-Ras in adult tissues to investigate its specificity. We discovered that the ability of K-Ras^G12D^ to influence the fitness of cells in a given tissue is not determined by its canonical signaling through MAPK. Instead, low baseline expression of c-Myc renders tissues non-permissive to oncogenic K-Ras, a context that can be reversed in the liver by ectopically expressing c-Myc. This functions independently of the proliferative index of the tissue or the induction of cell cycle arrest or apoptosis. Our findings reveal the importance of the basal state of the tissue-inherent signaling network for determining oncogene specificity.

## INTRODUCTION

Genomic profiling of primary cancers has revealed hundreds of genes that, when mutated, contribute to tumorigenesis. Nevertheless, mutational profiles of the majority of cancer drivers are not uniform across tumor types, but instead display remarkable selectivity for specific tissues-of-origin^1,2^. *KRAS* is a prime example of this mutational tropism. K-Ras is a member of the RAS protein family of small GTPases that transduce signals emanating from cell surface receptors to their effectors. Activating mutations in K-Ras promote the GTP-bound form of the protein, leading to constitutive downstream signaling that drives proliferation and survival^3^. *KRAS* mutations are only common in adenocarcinomas of the pancreas, colorectum, lung, endometrium, and biliary tract, as well as in multiple myelomas, but are exceedingly rare in cancers originating from other tissues, such as breast, prostate, brain, liver, and kidney^4^. It is unknown whether there is a tissue resident mechanism that actively promotes sensitivity or resistance to the pro-oncogenic signal emanating from mutant K-Ras. Understanding the molecular basis of tissue selectivity will lead to key insights into molecular determinants of oncogene activity.

Many factors could contribute to the phenomenon of tissue specificity, including the activity of mutational and DNA repair processes and the tissue-specific expression patterns of causal genes and/or functionally redundant proteins (in the case of tumor suppressors)^2,5^. Yet, these mechanisms provide an incomplete explanation for the specificity of most cancer drivers. For example, while the greater expression levels of K-Ras may underlie its heightened mutational frequency over its paralogs N-Ras and H-Ras^6^, the expressional profile of K-Ras itself does not correlate with its mutational frequency across tissues^7^. A critical determinant of tissue specificity may instead lie in the unique requirements of any genetic alteration for a permissive tissue context in order to exert its tumorigenic effect^2,8^. In support of this hypothesis, germline mutations in tumor suppressors that form the genetic basis of cancer predisposition syndromes manifest in the development of tumors in some, but not all tissues^9,10^.

In this study, we performed a systematic, comprehensive, and quantitative analysis of the oncogenic activity of K-Ras^G12D^ across ten tissues to provide unprecedented cellular and molecular resolution on the mechanisms of oncogene permissivity. We demonstrate that K-Ras permissivity is uncoupled from activation of the MAPK pathway, the basal proliferative rate of a tissue, and the induction of apoptosis and cell cycle arrest by K-Ras. Instead, we identified baseline expression of c-Myc as a molecular determinant of K-Ras permissivity. Furthermore, we show that elevating c-Myc levels is sufficient to establish a K-Ras permissive environment in the liver, an otherwise K-Ras resistant tissue. In general terms, our findings establish that the basal tissue-specific protein signaling network fundamentally determines the capacity of oncogenes to exert their pro-tumorigenic potential.

## RESULTS

### Modeling K-Ras tropism in mice

To identify the mechanisms of tissue-specific K-Ras oncogenicity, we first established a system for the expression of oncogenic K-Ras across multiple tissues in the adult animal, thus allowing us to experimentally define permissivity. We utilized the conditional *Kras*-LoxP-STOP-LoxP-G12D (*Kras*^LSL-G12D^) allele^11^ and evaluated two models expressing tamoxifen-inducible Cre recombinase (CreER) from promoters that are active in a wide range of tissues: *ROSA26*^Cre-ERT2^ and UBC-Cre-ERT2^TG^ ^12,13^. Both lines were crossed with mice carrying a recombination reporter (*ROSA26*^LSL-EYFP^ ^14^) to produce *ROSA26*^Cre-ERT2/LSL-EYFP^ (R26-CreER/YFP) and UBC-Cre-ERT2^TG^; *ROSA26*^LSL-EYFP/+^ (UBC-CreER;R26-YFP) mice. We assessed recombination efficiency by quantifying YFP^+^ cells in tissues of tamoxifen-treated and untreated control animals and found that UBC-CreER was more efficient and less leaky than R26-CreER across a range of tissues (Figure S1A and S1B). Heart was an outlier in this study. Recombination efficiency in the heart was poor in both models (Figure S1C); we thus expanded our assessment to a model in which the cardiac-specific α myosin heavy chain (*Myh6*) promoter controls expression of a tamoxifen-inducible Cre recombinase (MerCreMer)^15^. Efficient recombination throughout the heart was observed in *Myh6*-MerCreMer^TG/+^; *ROSA26*^LSL-EYFP/+^ (Myh6-CreER;R26-YFP) mice after tamoxifen treatment (Figure S1C).

We then set out to define which tissues are permissive versus resistant to K-Ras oncogenicity. For this, we crossed Myh6-CreER;R26-YFP and UBC-CreER;R26-YFP mice with animals carrying the conditional oncogenic *Kras*^LSL-G12D^ allele. As expression of K-Ras^G12D^ results in acute doubling of the K-Ras gene dosage in our models, we generated a new strain (*Kras*^LSL-WT^) to recapitulate this allele dosage for WT K-Ras, to be used as a control. Mice were separated into short-term and long-term experimental cohorts that underwent distinct tamoxifen treatment regimens (Figure 1A).

**Figure 1.**
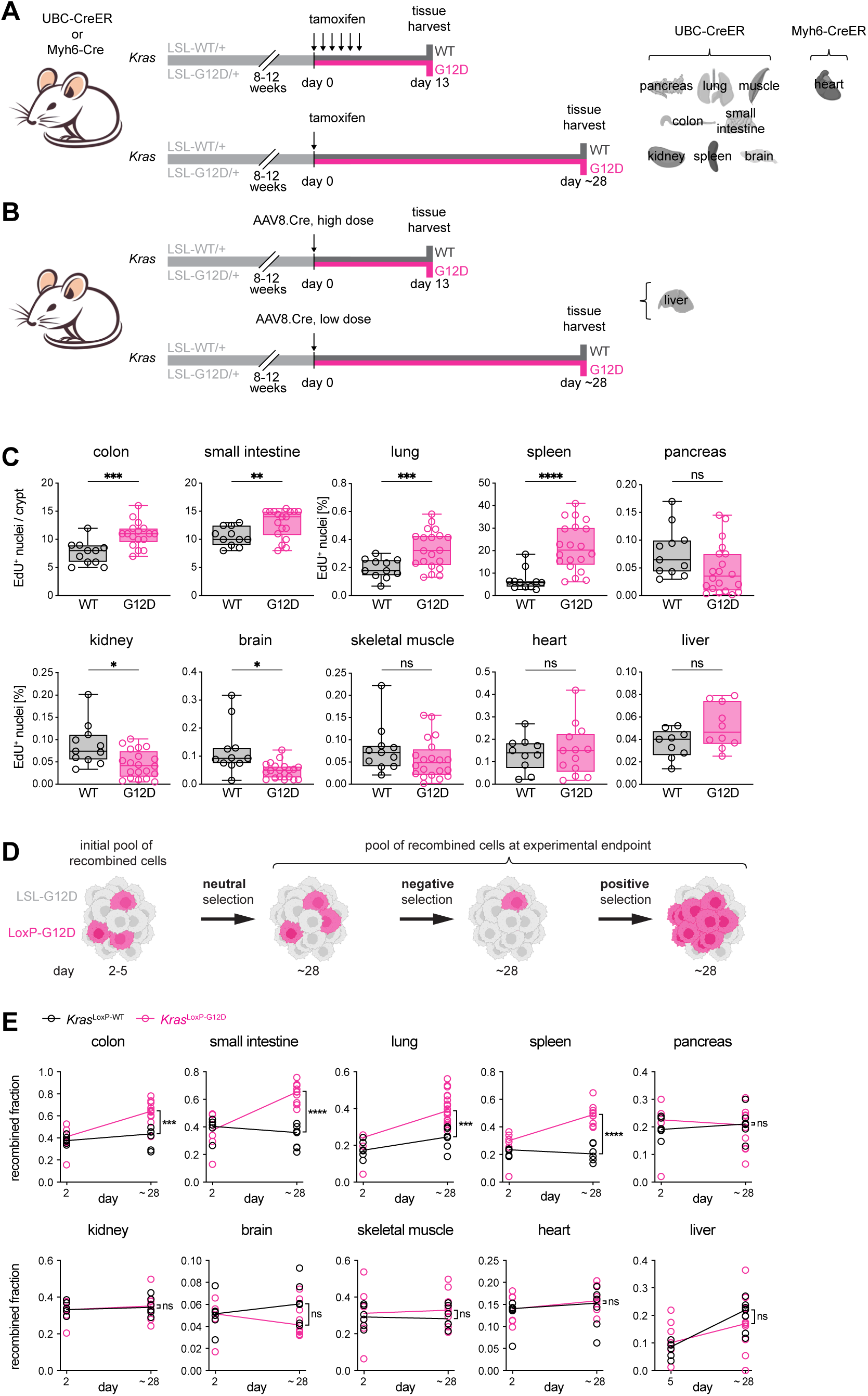
Expression of K-Ras^G12D^ confers an oncogenic phenotype only in a subset of tissues. (A) Experimental design for the investigation of the short-term (top) and long-term (bottom) effects of K-Ras^G12D^ expression across tissues. (B) Experimental design for the investigation of the short-term (top) and long-term (bottom) effects of K-Ras^G12D^ expression in the liver using viral delivery of Cre recombinase. (C) Cell proliferation measured by EdU incorporation across tissues from Myh6-CreER;R26-YFP (heart), R26-YFP (liver), and UBC-CreER;R26-YFP (all other tissues) mice carrying the conditional *Kras*-LSL-WT (K-Ras^WT^) or LSL-G12D (K-Ras^G12D^) allele treated with AAV8.Cre (liver) or tamoxifen (all other tissues) and harvested after ∼28 days. Plotted is the median number of EdU^+^ nuclei per crypt (colon, small intestine; min. 50 crypts per mouse) or percentage of EdU^+^ nuclei (all other tissues). Box plots indicate 25^th^ to 75^th^ percentile and group median. Statistical significance between group means was determined by Welch’s t-test. **** P < 0.0001, *** P < 0.001, ** P < 0.01, * P < 0.05, ns P > 0.05. (D) Concept behind the *in vivo* competition assay. The fraction of recombined, K-Ras^G12D^-expressing cells (magenta) changes over time depending on the tissue-specific effects of K-Ras^G12D^ on cellular fitness. (E) Fraction of cells with the recombined *Kras*-LSL-WT (*Kras*^LoxP-WT^, black) and *Kras*-LSL-G12D (*Kras*^LoxP-G12D^, magenta) allele at indicated timepoints, same experimental cohort as in (C). Lines connect group medians. Statistical difference in mean fraction of *Kras*^LoxP-WT^ vs. *Kras*^LoxP-G12D^ cells at day ∼28 was determined by Welch’s t-test. **** P < 0.0001, *** P < 0.001, ns P > 0.05. (A, B, and D) Figures partially created in BioRender. Popow, O. (2025) https://BioRender.com/jf1078h. See also Figure S1 and S2.

K-Ras^G12D^ protein was expressed in all evaluated tissues of short-term tamoxifen-treated UBC-CreER mice and the hearts of short-term tamoxifen-treated Myh6-CreER animals carrying the conditional oncogenic allele (Figure S1D). K-Ras^G12D^ expression translated into a detectable increase in levels of GTP-bound, *i.e.* active, Ras in all tissues, except the heart (Figure S1E). Examination of hematoxylin and eosin (H&E)-stained tissue sections from mice harvested one week after consecutive tamoxifen dosage revealed signs of inflammation in multiple tissues, most notably the spleen and liver, and this was independent of *Kras* genotype. Anti-CD45^+^ staining further confirmed that livers of tamoxifen-treated mice were infiltrated with leucocytes (Figure S2A). To disentangle treatment and liver-extrinsic effects, we utilized AAV8 carrying Cre recombinase expressed from the hepatocyte-specific thyroxide binding globulin (TBG) promoter (AAV8.TBG.PI.Cre.rBG/AAV8.Cre) to induce recombination exclusively in the liver (Figure 1B). Intravenous (IV) injection with AAV8.Cre resulted in expression of active K-Ras^G12D^ with no evidence of inflammation (Figure S2B-D).

In summary, we generated an array of GEMMs to systematically define K-Ras permissive versus K-Ras resistant tissues, creating an experimental system to uncover molecular mechanisms of K-Ras tissue specificity.

### K-Ras^G12D^ confers an oncogenic phenotype only in a subset of tissues

With our GEMM-based system in place to define the tissue-specific oncogenic activity of K-Ras, we first evaluated the histologic impact of K-Ras^G12D^ across the ten tissues of interest. K-Ras^G12D^-expressing mice exhibited colonic crypt hyperplasia and multifocal bronchioloalveolar hyperplastic lesions. We observed these effects in both short- and long-term experiments, with a more pronounced phenotype in the latter (Figure S2E). Treatment-associated inflammation in the spleen was absent in the long-term experiment, unmasking a pronounced K-Ras^G12D^-dependent effect in the form of increased extramedullary hematopoiesis (EMH) indicative of myeloproliferative disease (Figure S2E). Notably, the livers of these mice were also marked by EMH with mild to moderate lymphocytic infiltrates (Figure S2F). In contrast, livers of K-Ras^LSL-G12D^ animals in which recombination was induced by AAV8.Cre appeared histologically normal (Figure S2B, G), suggesting that the effect observed in the UBC-CreER model is liver/hepatocyte-extrinsic. No obvious histological changes were noted in response to K-Ras^G12D^ expression in the small intestine, pancreas, kidney, brain, skeletal muscle, or heart at either time point (Figure S2E and data not shown). Thus, K-Ras^G12D^ fails to produce a histological phenotype indicative of oncogenic activity in most of the assessed tissues.

To determine the effect of K-Ras^G12D^ on cell proliferation in a quantitative manner, we measured incorporation of the nucleoside analog 5-ethynyl 2’-deoxyuridine (EdU) into newly synthesized DNA ^16^. In line with our histological findings, expression of K-Ras^G12D^ resulted in increased proliferation in the lung, spleen, colon, and, notably, also the small intestine. K-Ras^G12D^ expression reduced proliferation in the kidney and brain. A similar trend was seen in the pancreas, although this was not significant (p = 0.075, Welch’s t test). We did not detect any significant differences in proliferation in the skeletal muscle, heart, or liver between K-Ras^WT^ and K-Ras^G12D^-expressing tissues (Figure 1C). Within the short-term experimental cohort, we observed a K-Ras^G12D^-dependent increase in proliferation only in the small intestine. Independent of *Kras* mutation status, tamoxifen treatment elicited elevated proliferation in the spleen, liver, skeletal muscle, and heart (Figure S2H). Proliferation was not altered in the livers of AAV8.Cre-treated animals of any *Kras* genotype (Figure S2I). Thus, K-Ras^G12D^-dependent induction of proliferation roughly correlates with histologic changes that occur in adult tissues when K-Ras^G12D^ is expressed.

We next asked whether K-Ras^G12D^ conferred increased cellular fitness in a tissue specific manner. We performed an *in vivo* cellular competition assay and used a droplet digital PCR (ddPCR)-based approach^17^ to measure the fraction of recombined (*Kras*^LoxP-G12D^ or *Kras*^LoxP-WT^) cells over time, which we reasoned would be reflective of cell fitness (Figure 1D). To establish the initial pool of recombined cells, we quantified the fraction of *Kras*^LoxP^ cells two days after tamoxifen and five days after AAV8.Cre dosage. In addition, we determined the fraction of recombined cells after ∼28 days. We observed that the percentage of *Kras*^LoxP-WT/+^ cells remained relatively constant in all assayed tissues from tamoxifen-treated UBC-CreER and Myh6-CreER animals, indicating that doubling the dosage of WT K-Ras does not provide a fitness advantage *in vivo* in any tissue (Figure 1E). Notably, *Kras*^LoxP-G12D/+^ cells were not eliminated, but instead remained present at fixed proportions in the pancreas, kidney, brain, skeletal muscle, and heart (Figure 1E); tissues that were classified as non-permissive based on histology and proliferation (Figure S2E and 1C). In contrast, the fraction of *Kras*^LoxP-G12D^ cells increased over time in permissive tissues (colon, small intestine, lung, and spleen; Figure 1E) and this was in line with the positive effect of K-Ras^G12D^ expression on proliferation in these tissues (Figure 1C). In the livers of AAV8.Cre treated animals the fraction of *Kras*^LoxP-WT/+^ and *Kras*^LoxP-G12D/+^ cells increased over time, indicating ongoing recombination after measurement of the initial pool of *Kras*^LoxP^ cells after five days. Importantly, both cell populations increased at a similar rate, suggesting the absence of a competitive advantage of *Kras*^LoxP-G12D/+^ over *Kras*^LoxP-WT/+^ cells in the liver (Figure 1E).

Taken together, we established a system that allows us to experimentally define the tissues that are K-Ras permissive versus K-Ras resistant. Expression of K-Ras^G12D^ elicited a pro-tumorigenic response – measured as an effect on tissue histology, cell proliferation, and fitness – only in a subset of tissues, namely the spleen, small intestine, lung, and colon. In contrast, our three independent analyses categorize the heart, skeletal muscle, kidney, liver, pancreas, and brain as tissues that are non-permissive to the effects of oncogenic K-Ras expression (Figure S2J).

### K-Ras permissive tissues are characterized by a pro-proliferative molecular signature

Oncogenic K-Ras can signal through a range of downstream effector pathways. The components of these signaling cascades are expressed at variable levels across tissues. We thus investigated whether specific differences in a tissue’s steady-state signaling network will determine its response to an oncogenic stimulus, such as the expression of K-Ras^G12D^. To this end, we applied global, untargeted Tandem Mass Tag (TMT)-based quantitative mass spectrometry (MS) to profile the proteome of the ten tissues we included in this study. A total of 9,494 proteins were quantified in all channels of at least one TMTplex (Table S1). Replicate samples clustered together in principal component analysis (PCA), indicating good reproducibility across plexes (Figure S3A).

To identify molecular features that distinguish K-Ras permissive from non-permissive tissues, we first correlated the protein log_2_ sum intensity with the K-Ras^G12D^ fitness score (Figure S2J) across tissues to obtain a Spearman rank coefficient estimate for each protein in our dataset (Figure S3B). This analysis ranked proteins based on the association between their expression across tissues and the K-Ras permissivity of these same tissues. We then performed Gene Set Enrichment Analysis (GSEA) against this ranked list of proteins using the MSigDB mouse-ortholog hallmark gene set collection (Figure 2A, Figure S3C). Twenty gene sets were significantly enriched (normalized enrichment score (NES) > 1.3; −log_10_(FDR) > 1) among the proteins positively associated with K-Ras permissivity and five gene sets were significantly enriched (NES < −1.3; −log_10_(FDR) > 1) among the proteins negatively associated with K-Ras permissivity. Notably, many of the positively enriched gene sets relate to pathways or cellular processes involved in cell proliferation and inflammation. We further performed a leading edge analysis to identify the proteins that contributed to the enrichment of pathways associated with proliferation and the overlap between them (Figure 2B). Only the ‘E2F Targets’ and ‘G2M Checkpoint’ gene sets shared a considerable proportion of leading edge proteins. In contrast, the enrichment of all other proliferation-related gene sets was driven by a distinct set of proteins. A total of 299 leading edge proteins were unique to a single gene set.

**Figure 2.**
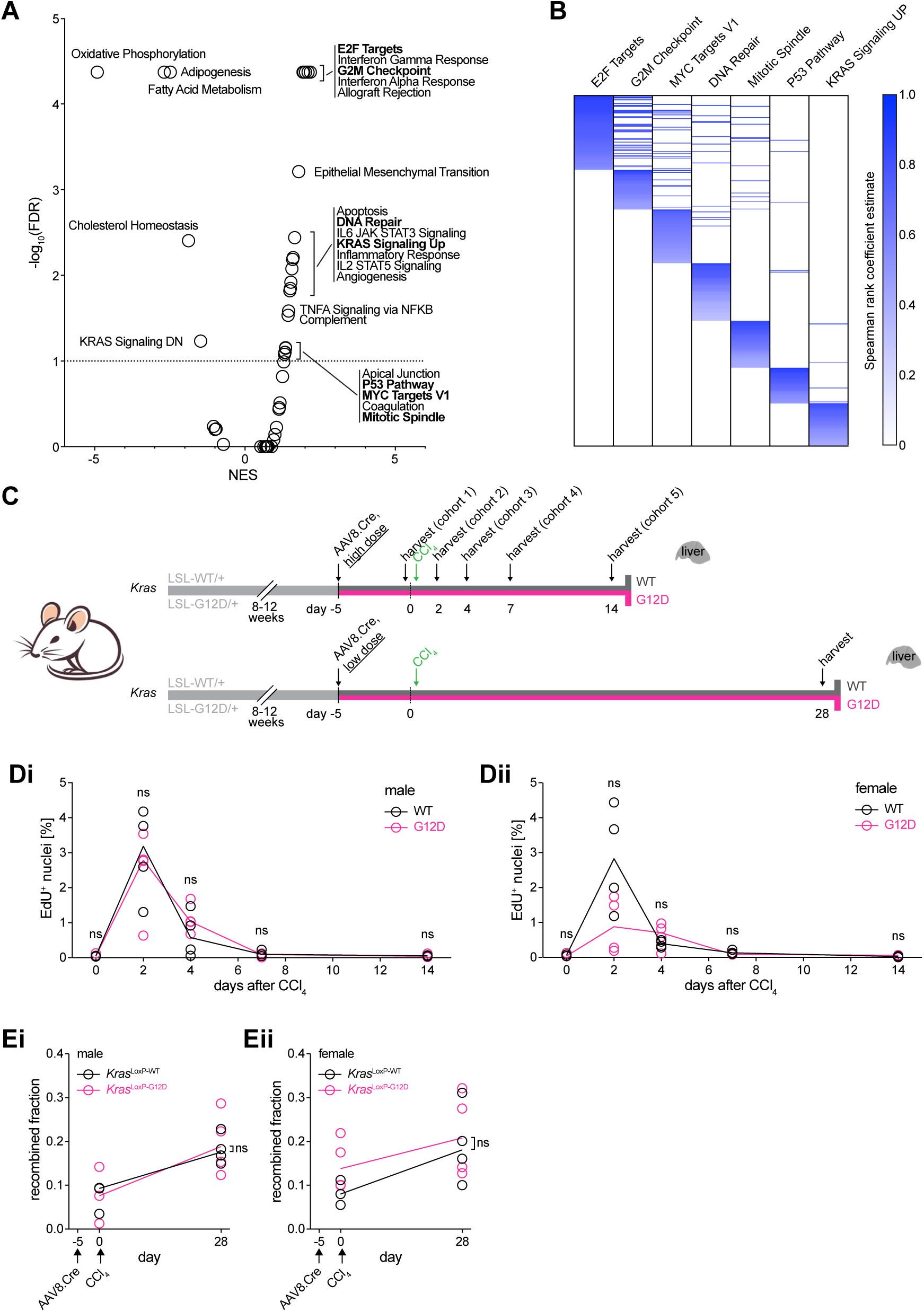
An acute increase in proliferation is not sufficient to create a K-Ras^G12D^-permissive environment. (A) Enrichment of hallmark pathway gene sets relative to K-Ras^G12D^ permissivity as determined by GSEAPreranked. Plotted is the normalized enrichment score (NES) against the −log_10_ false discovery rate (FDR), horizontal dashed line indicates the significance cut-off at −log_10_(FDR) ≥ 1. Names of significantly enriched pathways broadly related to cell proliferation are highlighted in bold. (B) Overlap in leading edge proteins between hallmark gene sets highlighted in (A). Proteins are colored based on their correlation coefficient between relative expression and K-Ras permissivity (Figure S3B). (C) Experimental design to assess the oncogenicity of K-Ras^G12D^ in the presence of proliferation in the liver. (D) Male (i) and female (ii) R26-YFP mice carrying the conditional *Kras*-LSL-WT (WT, black) or LSL-G12D (G12D, magenta) allele injected with 2×10^11^ GCs of AAV8.Cre (day −5) and CCL_4_ (day 0) and harvested at indicated days after CCL_4_ administration. Animals harvested at day 0 did not receive CCL_4_. Plotted is the percentage of EdU^+^ nuclei. (E) Recombined fraction of the *Kras*^LoxP-WT^ (WT, black) and *Kras*^LoxP-G12D^ (G12D, magenta) allele in the livers of R26-YFP mice injected with 0.5×10^10^ (male, Ci) or 1×10^10^ (female, Cii) GCs of AAV8.Cre on day −5 and harvested five days later (day 0, no CCL_4_), or 28 days after CCL_4_ administration. The increase in the recombined fraction between day 5 and 28 is likely attributable to a continuation of AAV8.Cre-mediated recombination beyond day 5 (see also Figure 1E). (D and E) Lines connect group medians. Statistical significance between group means was determined by Welch’s t-test. Ns P > 0.05. See also Figure S3.

These findings indicate that, at steady state, K-Ras permissive tissues are characterized by a broad pro-proliferative/inflammatory proteomic signature encompassing hundreds of proteins. This is in line with our observation that K-Ras permissive tissues display an overall higher baseline proliferation rate compared to non-permissive tissues (Figure 1C).

### An acute increase in proliferation is not sufficient to create a K-Ras^G12D^-permissive environment

Our proteomic analysis suggested that the homeostatic proliferative state of a tissue might be predictive of its permissivity to K-Ras^G12D^. We therefore tested whether an increase in cell proliferation is sufficient to create a K-Ras permissive tissue environment. We chose the liver as an exemplar of a non-permissive tissue, as its low levels of baseline proliferation are highly amenable to manipulation with carbon tetrachloride (CCL_4_). CCL_4_ causes acute liver injury, triggering an inflammatory response and tissue regeneration through a temporary increase in hepatocyte proliferation^18,19^. We first induced K-Ras^G12D^ expression in the livers of mice and then injected them with CCL_4_ (Figure 2C). We separately analyzed livers from male and female mice, as sex differences in the response to CCL_4_ have previously been reported^20^. CCL_4_ treatment resulted in hepatocyte necrosis and immune cell infiltration that completely resolved within seven days (Figure S3D). Concomitantly, cell proliferation increased dramatically, peaking two days after CCL_4_ administration, and subsequently returned to baseline levels by day 14 (Figure 4B). We did not detect any significant differences in the histological or proliferative responses between livers expressing K-Ras^WT^ or K-Ras^G12D^ from male or female mice.

We next assessed whether K-Ras-mutant hepatocytes preferentially accumulate over wild-type cells during tissue regeneration. For this experiment, we used mice of the same genotypes, but injected a lower dose of AAV8.Cre to induce recombination in only ∼10% of hepatocytes prior to treatment with CCL_4_ (Figure 2C, bottom panel). We did not detect any significant difference in the mean fraction of *Kras*^LoxP-WT^ vs. *Kras*^LoxP-G12D^ cells at endpoint as determined by ddPCR (Figure 4C). Taken together, our results demonstrate that a temporary boost in the proliferative index and the presence of an inflammatory environment are not sufficient to convert the liver into a K-Ras-permissive tissue.

### Cell cycle arrest or apoptosis are not universal resistance mechanisms to K-Ras^G12D^ expression in non-permissive tissues

We next considered an alternative hypothesis: that an active resistance mechanism was functioning in tissues that were not permissive to K-Ras^G12D^. Mutant forms of Ras have been reported to induce senescence in premalignant lesions^21,22^ and to engage pro-apoptotic effector pathways^23–,25^. Moreover, in the absence of cooperating co-mutations, ectopic expression of oncogenic Ras results in a p19^Arf^-dependent cell cycle arrest, and is accompanied by increased expression of the CDK inhibitor p21^Cip1/Waf^ ^26–,28^. To address the potential for negative homeostatic effects of oncogenic K-Ras, we first quantified p21^Cip1/Waf^ protein expression in tissues after acute induction of K-Ras^G12D^. Levels of p21^Cip1/Waf^ significantly increased in a K-Ras^G12D^-dependent manner in the small intestine, colon and skeletal muscle. No significant changes were observed in the other tissues (Figure 3A, Figure S4A). Next, we detected apoptotic cells by terminal deoxynucleotidyl transferase-dUTP nick end labeling (TUNEL) followed by fluorescence microscopy. The percentage of TUNEL^+^ cells was low across all experimental conditions with a median value below 1%. K-Ras^G12D^ expression did not result in increased apoptosis in any of the assayed tissues (Figure 3B). Collectively these results indicate that neither cell cycle arrest nor apoptosis are general resistance mechanisms of K-Ras non-permissive tissues.

**Figure 3.**
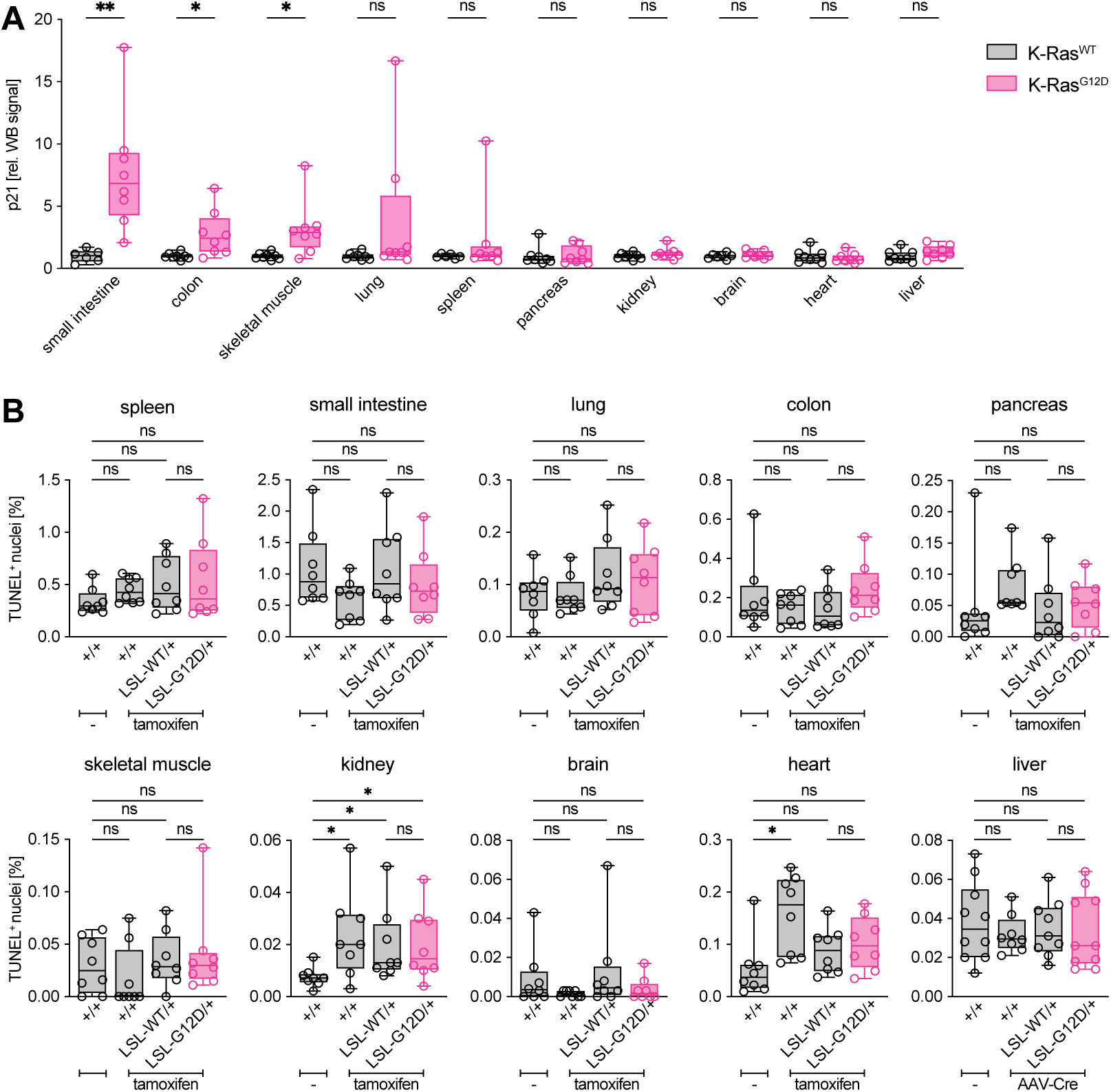
Cell cycle arrest or apoptosis are not universal resistance mechanisms to K-Ras^G12D^ expression in non-permissive tissues. (A) Protein expression of p21^Cip1/Waf^ in tissue lysates from R26-YFP (liver), Myh6-CreER;R26-YFP (heart), and UBC-CreER;R26-YFP (all other tissues) mice carrying a conditional wild-type (*Kras*^LSL-WT^) or G12D (*Kras*^LSL-G12D^) *Kras* allele injected with 2×10^11^ GCs of AAV8.Cre and harvested 13 days after (liver) or treated on six consecutive days with tamoxifen and harvested seven days after (all other tissues). Plotted is the relative anti-p21^Cip1/Waf^ WB signal normalized to vinculin. Statistically significant differences between group means were determined by Welch’s t-test. (B) Apoptotic cells quantified by TUNEL and fluorescence microscopy in tissue sections from the same experimental cohort as in (A) but also including mice without a conditional *Kras* allele (+/+). Statistically significant differences between means were determined by Brown-Forsythe and Welch ANOVA tests. (A and B) Box plots indicate 25^th^ to 75^th^ percentile and group median. ** P < 0.01, * P < 0.05, ns P > 0.05. See also Figure S4.

### Members of the RAS pathway are collectively expressed at higher levels in K-Ras permissive tissues

We next focused our analysis of baseline differences between K-Ras permissive and non-permissive tissues on the more immediate protein signaling network surrounding Ras. Because low abundant proteins – such as transcription factors – often evade reproducible quantification by shotgun proteomics, we instead applied TOMAHAQ (triggered by offset, multiplexed, accurate-mass, high-resolution, and absolute quantification)/Tomahto – a targeted TMT-based approach^29,30^. We identified 482 suitable trigger peptides encompassing 186 murine members of the RAS pathway^31^; these were unique and cysteine-free peptides lacking trypsin miscleavage sites that had previously been detected by MS-based proteomics in any of three large scale studies^32–,34^. Because TOMAHAQ/Tomahto does not require sample fractionation, instrument time is dramatically reduced – allowing us to analyze protein expression across tissues from individual mice, instead of pooling samples as before. The mixture of 482 trigger peptides labeled with super heavy TMT was spiked into the samples containing the endogenous TMT1-10-labeled tissue peptides and the TMT11-labeled bridge channel. These were then analyzed following the TOMAHAQ/Tomahto workflow described previously^30^.

Out of 482 trigger peptides, 467 were detected in at least one sample (mean = 440/plex) and these corresponded to 184 proteins (mean = 182/plex, Figure S5A). At the endogenous level, we identified 418 (mean = 325/plex) and quantified 384 peptides (mean = 279/plex) in at least one sample (Figure S5Bi). These translated to 176 identified (mean = 158/plex) and 170 quantified proteins (mean = 148/plex; Figure S5Bii, Table S2). PCA demonstrated good reproducibility across samples (Figure S5C). Out of all quantified proteins, the epidermal growth factor receptor was the only one to display significant differences in expression between male and female mice across several tissues, with males consistently expressing higher levels (Figure S5D).

We assessed how K-Ras tissue specificity related to RAS pathway protein expression by calculating the Spearman rank correlation coefficients between the protein log_2_ sum intensities of each quantified member of the RAS pathway with the K-Ras^G12D^ fitness score across tissues. Strikingly, the vast majority of pathway members – including negative regulators – displayed positive correlation coefficients, *i.e.* were expressed at higher levels in K-Ras permissive compared to non-permissive tissues (Figure 4). Consistent with prior work^6^, protein expression of K-Ras or its paralogues did not significantly correlate with K-Ras permissivity. A closer look at the two splice isoforms revealed that while K-Ras4b was most abundant in the brain, followed by lung and spleen; K-Ras4a was expressed at higher levels in the colon and small intestine compared to the other tissues (Figure S5E). The Mitogen-activated protein kinase (MAPK) pathway is a central axis of K-Ras signaling, mediating many of its pro-oncogenic effects^3^. Our targeted MS-based assay revealed that the expression of c-Raf (Raf-1) and extracellular signal-regulated kinase 1 (ERK1/MAPK3) were significantly and positively correlated with a tissue’s permissivity to K-Ras^G12D^ (Figure 4).

**Figure 4.**
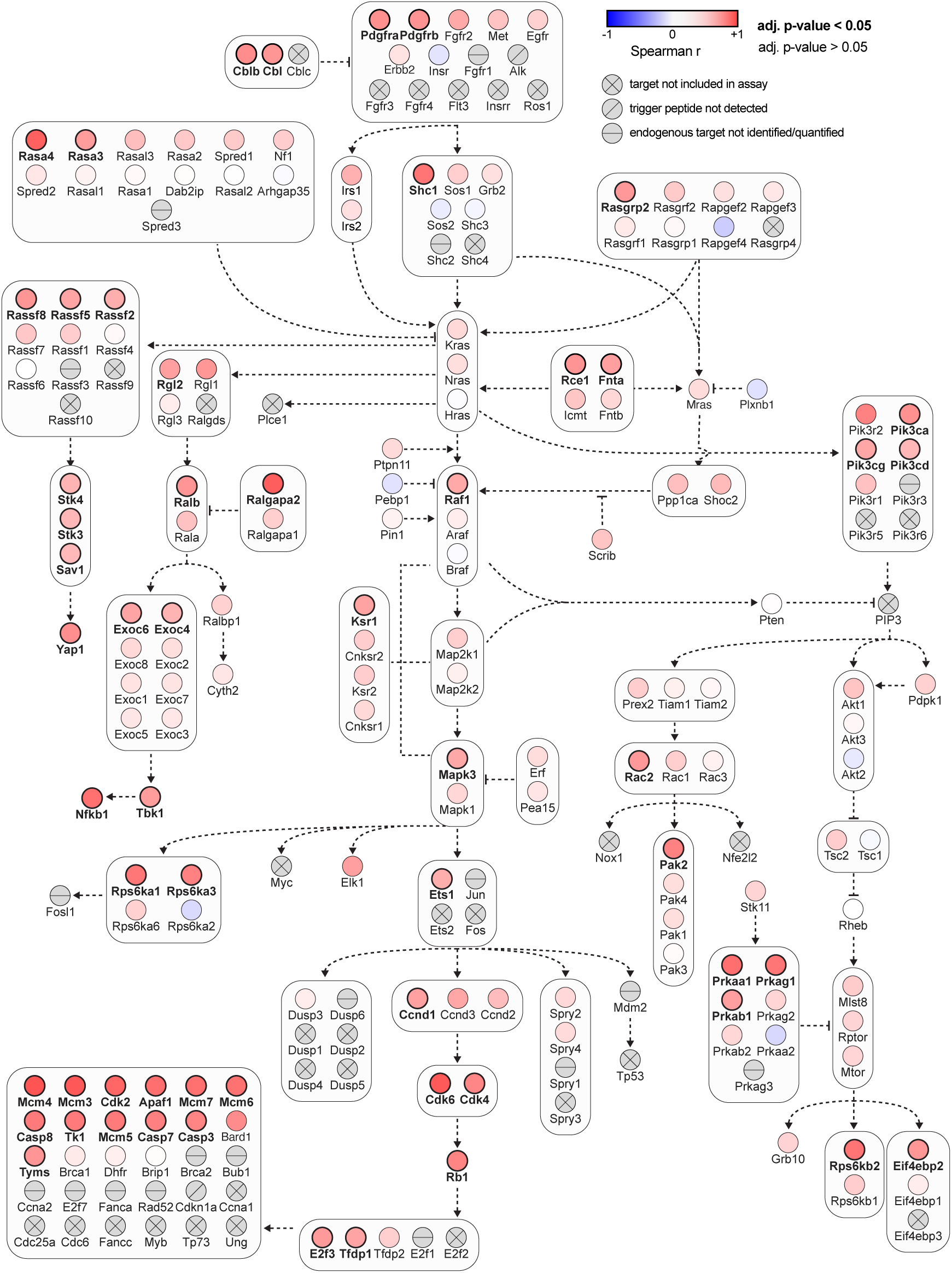
RAS pathway components are collectively expressed at higher levels in K-Ras permissive tissues. RAS pathway diagram adopted from ^31^, each member is labeled with its murine gene name. Coloring of nodes indicates the degree and directionality of the correlation between relative protein expression (measured by TOMAHAQ/Tomahto) and tissue permissivity to K-Ras^G12D^. Positive correlations (Spearman r = 0-1) are shown in red and negative correlations (Spearman r = −1-0) in blue. P values were adjusted for multiple testing using Bonferroni correction, proteins with an adj. p value < 0.05 are highlighted in bold. Proteins which were not quantified are shown in grey: proteins that were not included in the assay because no suitable trigger peptide could be identified are indicated with a cross; proteins which were included in the assay but for which its corresponding trigger peptide(s) were not identified are indicated with a slanting line; proteins for which its corresponding endogenous peptide(s) were not identified or quantified in all channels of at least one TMTplex are indicated with a horizontal line. See also Figure S5.

In summary, our results indicate that no single protein, including K-Ras itself, or group of proteins within the RAS pathway distinguish K-Ras permissive from non-permissive tissues. Instead, the collective K-Ras signaling network is elevated in permissive tissues.

### MAPK activation is not sufficient to mediate K-Ras permissivity

As the MAPK pathway is a central axis of K-Ras signaling, and we observed up-regulated c-Raf and ERK1 in permissive tissues (Figure 4), we tested whether the ability of K-Ras^G12D^ to activate the MAPK pathway was restricted to permissive tissues. We utilized an enzyme-linked immunosorbent assay (ELISA) to measure levels of total ERK1/2 and active ERK1/2 phosphorylated at Thr202/Tyr204. Expression of K-Ras^G12D^ induced ERK phosphorylation in most tissues and this was independent of their K-Ras permissivity (Figure 5A). We did not detect a significant increase in ERK activation in the lung, consistent with the absence of phosphorylated ERK from K-Ras^G12D^-dependent low-grade lung lesions^35^. ERK1/2 expression levels were low in both skeletal and heart muscle and fell below the detection limit of the ELISA. We therefore assessed ERK phosphorylation in these tissues by Western blotting (WB, Figure S6A). Expression of K-Ras^G12D^ resulted in increased ERK phosphorylation in the skeletal muscle, which predominantly expressed ERK2 (Figure S6Ai). We did not observe any changes in ERK activation in response to K-Ras^G12D^ expression in the heart and this was in line with the lack of a measurable increase in Ras-GTP levels in this tissue (Figure S1E).

**Figure 5.**
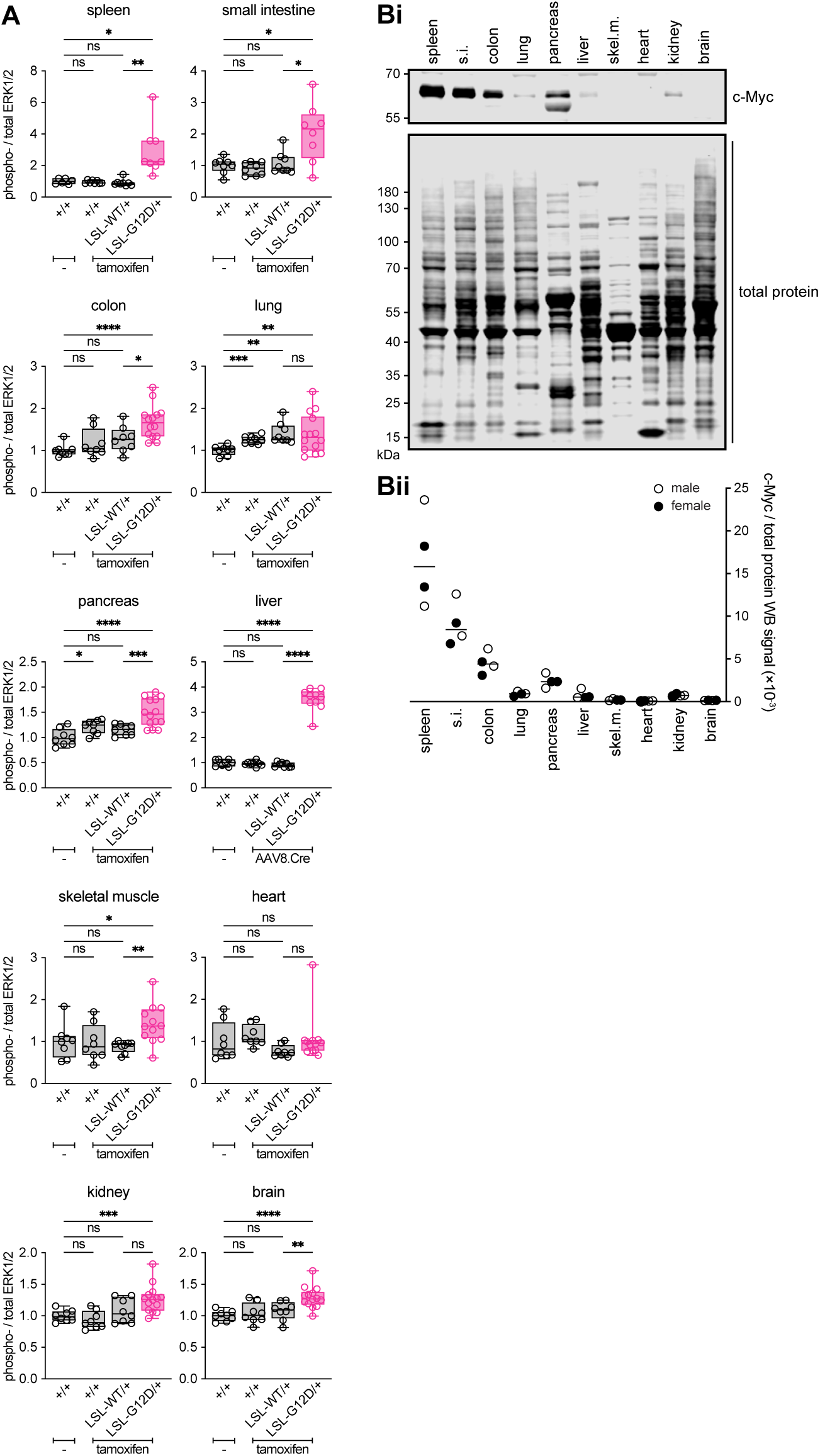
MAPK activation is not sufficient to mediate K-Ras permissivity. (A) Abundance of ERK1/2 phosphorylated at Thr202/Tyr204 (phospho-ERK) relative to total ERK1/2, determined by WB (skeletal muscle and heart) or ELISA (all other tissues). Samples were from R26-YFP (liver), Myh6-CreER;R26-YFP (heart), and UBC-CreER;R26-YFP (all other tissues) mice carrying the indicated *Kras* alleles and injected with 2×10^11^ GCs of AAV8.Cre and harvested 13 days after (liver) or treated on six consecutive days with tamoxifen and harvested seven days after (all other tissues). Box plots indicate 25^th^ to 75^th^ percentile and group median. Statistically significant differences between means were determined by Brown-Forsythe and Welch ANOVA tests. **** P < 0.0001, *** P < 0.001, ** P < 0.01, * P < 0.05, ns P > 0.05. (B) Expression of c-Myc across wild-type tissues. (Bi) Representative blot of tissues harvested from a female mouse. (Bii) Aggregated measurements across four individual mice; males are indicated as empty, females as filled circles. Plotted is the relative c-Myc abundance relative to total protein signal. See also Figure S6.

Translocation of phosphorylated ERK from the cytoplasm to the nucleus, where many of its substrates reside, is a critical step in the MAPK signaling pathway^36^. Indeed, the absence of nuclear phospho-ERK has been proposed as a potential mechanism of resistance to oncogenic K-Ras expression^37^. We evaluated the capacity of K-Ras^G12D^ to induce nuclear translocation of phosphorylated ERK1/2 by IF. Nuclear phospho-ERK1/2 was present in all tissues expressing K-Ras^G12D^ irrespective of their permissivity to K-Ras (Figure S6B). These results suggest that MAPK activation, including the translocation of phosphorylated ERK1/2 to the nucleus, is not sufficient to elicit an oncogenic response downstream of K-Ras^G12D^.

### c-Myc is differentially expressed between K-Ras permissive and non-permissive tissues

As canonical MAPK signaling activation did not correlate with K-Ras permissivity, we sought to test alternative hypotheses for the signaling network-based mechanism of K-Ras tissue specificity. Our shotgun proteomics experiment identified an enrichment for c-Myc targets in K-Ras permissive tissues (Figure 2A). c-Myc functions downstream of the MAPK pathway; phosphorylation of c-Myc at serine 62 by ERK inhibits its proteasomal degradation and allows for increased transcription of its target genes^38^. Unfortunately, because of its low abundance, c-Myc is conspicuously absent from most shotgun proteomic data sets, including the three studies we used to identify suitable trigger peptides for our targeted MS-based assay. Adapting a previously described protocol to prevent c-Myc degradation in cell lysates^39^, we used trichloroacetic acid (TCA) precipitation prior to protein resolubilization in WB loading buffer to evaluate endogenous c-Myc protein expression across tissues. With the exception of pancreas, all K-Ras non-permissive tissues displayed exceedingly low levels of c-Myc protein. In contrast, spleen, colon, and small intestine expressed high amounts of c-Myc relative to the other tissues (Figure 5B).

### Ectopic expression of c-Myc converts the liver into a K-Ras permissive tissue

Based on this observation, we reasoned that baseline c-Myc levels might be limiting for K-Ras permissivity and that, by extension, ectopic expression of c-Myc would be able to convert a K-Ras non-permissive to a permissive tissue. To test this hypothesis, we utilized the *ROSA26*^CAG-LSL-MYC/+^ GEMM^40^ and crossed it to mice carrying a conditional wild-type (R26-MYC;KrasWT) or mutant *Kras* allele (R26-MYC;KrasG12D). Induction of recombination in ∼100% of hepatocytes in R26-MYC;KrasWT mice by AAV8.Cre (Figure 6A) resulted in ectopic expression of c-Myc in the liver at levels comparable to those observed in the colon and small intestine (Figure S7A). Activation of ERK in the livers of AAV8.Cre-treated R26-MYC;KrasG12D mice was accompanied by an increase in both c-Myc phosphorylation at serine 62 and total c-Myc protein relative to R26-MYC;KrasWT livers (Figure S7B and S7C). This indicates that the post-translational regulation of c-Myc downstream of oncogenic K-Ras is maintained in this organ.

**Figure 6.**
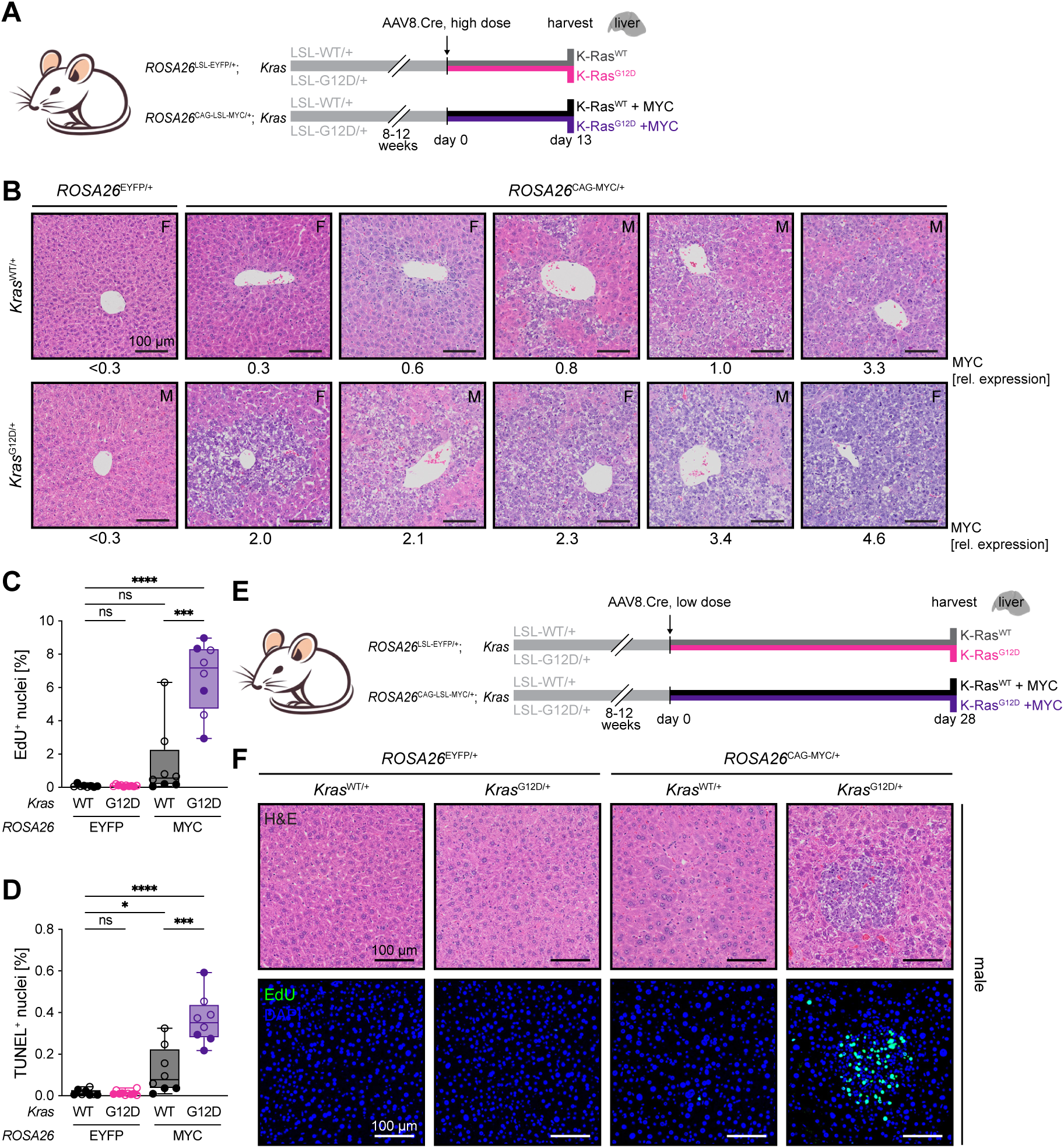
Ectopic expression of c-Myc creates a K-Ras permissive environment in the liver. (A) Experimental design to assess the short-term effects of K-Ras^G12D^ expression in the absence or presence of ectopic c-Myc. (B) Representative H&E-stained sections from mice ectopically expressing c-Myc and K-Ras^WT^ (top row) or K-Ras^G12D^ (bottom row) in ∼100% of hepatocytes. Liver sections from control animals expressing EYFP are shown on the left. Mice were harvested 13 days after treatment with 2×10^11^ GCs of AAV8.Cre. Relative c-Myc expression in the individual liver samples as determined in Figure S7Ciii is indicated below each image. Sex of animals is denoted in the top right corner of each image (M: male, F: female). Scale bars = 100 μm. (C and D) Cell proliferation measured by EdU incorporation (C) and apoptosis measured by TUNEL (D) in livers of mice ectopically expressing either EYFP or c-Myc and K-Ras^WT^ or K-Ras^G12D^. Same experimental cohort as in (B). Box plots indicate 25^th^ to 75^th^ percentile and group median. Samples from male mice are shown as empty circles, samples from female mice as filled circles. Statistically significant differences between means were determined by Brown-Forsythe and Welch ANOVA tests. **** P < 0.0001, *** P < 0.001, * P < 0.05, ns P > 0.05. (E) Experimental design to assess the long-term effects of induction of K-Ras^G12D^ expression in the absence or presence of ectopic c-Myc in a subset of hepatocytes. (F) Representative liver images from male mice ectopically expressing either EYFP or c-Myc and K-Ras^WT^ or K-Ras^G12D^ and harvested after 28 days. Proliferating cells are marked by anti-EdU staining. See also Figure S7.

Ectopic c-Myc expression induced widespread hepatocellular atypia, cytologically resembling carcinoma, within two weeks. The severity of this phenotype was variable across mice, but markedly correlated with relative c-Myc expression levels, which were highest in R26-MYC;KrasG12D animals (Figure 6B). Similarly, the degree of both proliferation and apoptosis induced by c-Myc strongly correlated with its expression levels (Figure S7D and S7E), consistent with previous findings^41^. Accordingly, livers expressing K-Ras^G12D^ and c-Myc were more proliferative compared to livers expressing c-Myc alone (Figure 6C). Notably, K-Ras^G12D^ expression did not inhibit c-Myc-induced apoptosis in this setting (Figure 6D).

To model a more physiologically relevant situation, we repeated this experiment, but induced recombination in only a subset of hepatocytes, and harvested livers after 28 days (Figure 6E). As noted above, expression of K-Ras^G12D^ alone did not elicit a phenotype in the liver. In agreement with a previous report^42^, ectopic expression of c-Myc induced nuclear enlargement; a phenotype that was more pronounced in male mice. Of the ten R26-MYC;KrasWT animals included in this experiment, only one male developed several small (<100 μm) tumor nodules. In contrast, expression of K-Ras^G12D^ in the context of ectopic c-Myc expression resulted in the formation of hepatocellular carcinomas in 10/10 animals. Additionally, the surrounding tissue was marked by moderate atypia characterized by nuclear enlargement and irregular chromatin. Once again, male mice exhibited a stronger phenotype including a higher number and larger size of lesions (Figure 6F and S7F). Tumor nodules in R26-MYC;KrasG12D mice were highly proliferative as indicated by increased EdU incorporation (Figure 6F and S7F). In summary, our results demonstrate that elevation of baseline c-Myc expression levels to those observed in permissive tissues is sufficient to create a K-Ras permissive environment in the liver, thus identifying c-Myc as a permissivity factor for oncogenic K-Ras.

## DISCUSSION

Although *KRAS* is a frequently mutated oncogene, the breath of its mutational profile is restricted to a small number of cancers. The mechanisms driving this tissue specificity are not well understood. In this work, we generate new mouse models to experimentally define tissues that are permissive or non-permissive to oncogenic K-Ras. We identify that colon, small intestine, lung, and spleen/blood are K-Ras permissive, while brain, heart, skeletal muscle, liver, kidney, and pancreas are K-Ras resistant. Furthermore, we demonstrate that K-Ras permissivity, defined in-part by its induction of proliferation, is uncoupled from apoptosis and cell cycle arrest. Using deep proteomic approaches we identify molecular signatures that correlate with K-Ras permissivity, ultimately leading us to c-Myc as a determinant of K-Ras permissivity, as demonstrated by our finding that ectopic c-Myc expression is sufficient to convert a K-Ras resistant tissue (liver) to become K-Ras permissive. In sum, we map the landscape of K-Ras permissivity at unprecedented resolution, providing new insights into mechanisms of tissue specificity in cancer.

Tissue specificity is a characteristic shared across many - if not most - cancers drivers. Oncogene specificity is generally solely defined by mutational profiles because studies that experimentally map the tissue specific oncogenic activity of cancer drivers are lacking. Ultimately, this limits our understanding of the molecular mechanisms underlying tissue specificity. As an example, *KRAS* mutations could be absent from a given tumor type for a variety of reasons, such as a lack of K-RAS expression or a neutral or tumor suppressive effect of mutant K-RAS in the cell-of-origin. Understanding the true susceptibility of a tissue to an oncogenic *KRAS* mutation thus necessitates an experimental model. The study presented here addresses this gap by systematically and quantitatively defining K-Ras tissue permissivity experimentally utilizing GEMMs.

We measured the effect of K-Ras^G12D^ expression on overall cell fitness and found that the hematopoietic system – represented by the spleen – was, by far, the most permissive to oncogenic K-Ras, followed by small intestine, lung, and colon. In contrast, the six other tissues we assessed – liver, kidney, pancreas, brain, skeletal muscle, and heart – were non-permissive. Our experimentally based classification largely aligns with the mutational frequency of *KRAS* in primary cancers. The inconsistencies, however, highlight the limitations of human data as an indirect predictor of tissue permissivity and also offer important insights. Our analysis clearly indicates that the majority of cells in the adult pancreas are non-permissive to oncogenic K-Ras, yet approximately 90% of pancreatic ductal adenocarcinomas (PDACs) carry *KRAS* mutations^4^. This observation is consistent with the idea that only a small sub-population of pancreatic cells – presumably representing the cells-of-origin of PDAC – is susceptible to oncogenic K-Ras expression. Moreover, the disease etiology of PDAC may play a decisive role in shaping K-Ras permissivity in this specific tissue context. Pancreatitis is a known risk factor for the development of PDAC^43^. Concordantly, inflammation and regeneration after injury can sensitize pancreatic cells to the effects of oncogenic K-Ras in mouse models^44^. This suggests that non-permissive environments can be altered to create permissive cellular contexts. We show that CCL_4_-induced inflammation and compensatory proliferation are, however, not sufficient to create a K-Ras permissive environment in the liver. Although we only tested the effects of an acute inflammatory insult, human data indicate that chronic inflammation, for example in the context of liver cirrhosis, is also not associated with *KRAS*-mutant hepatocellular carcinoma^45,46^.

Here, we assessed the permissivity of adult tissues to K-Ras^G12D^, the most frequent allele in human cancers. A change in these parameters will undoubtedly affect K-Ras tissue specificity. First, the permissivity of a tissue can shift in the course of development. For example, the embryonic heart – in contrast to its adult counterpart as shown here – is sensitive to mutant K-Ras expression^47,48^. Likewise, the capacity of oncogenic K-Ras to induce pancreatic intraepithelial neoplasia (PanIN) formation drastically reduces as the pancreas matures^44,49^. Second, tissue permissivity varies between mutant alleles. This is illustrated by the distinct *KRAS*^Mut^ allele frequencies observed in cancers originating from different tissues. Importantly, these differences can only be in part explained by tissue-specific mutational processes^50^. As an example, K-Ras^G13D^ promotes hyperproliferation of colonic crypts^51^ but is – unlike K-Ras^G12D^ – unable to robustly induce adenoma formation in the lung^52^. Consistently, *KRAS*^G13D^ mutations are enriched in colorectal cancers, but absent from lung tumors^50^. Lastly, we assessed K-Ras permissivity in the absence of any co-mutations. While it is conceivable that the loss of a potent tumor suppressor, such as p53, would be sufficient to overcome resistance to mutant K-Ras, existing data argue against this hypothesis. The tumor spectrum in mice that spontaneously activate K-Ras in all tissues did not significantly differ between p53 proficient and deficient backgrounds. Indeed, the only additional tumor type that emerged in the absence of p53 were sarcomas^53^. Consistently, deletion of the *Arf* locus, which encodes the p53-stabilizing protein p19^Arf^, also allowed for sarcoma formation in this model, without altering the remaining tumor spectrum^54^. This is in line with our observation that the skeletal muscle is the only non-permissive tissue in which the CDK inhibitor and p53 target gene p21^Cip1/Waf^ is upregulated in response to oncogenic K-Ras expression. Despite this example, our study indicates that selection against expression of oncogenic K-Ras is not a common mechanism underlying its tissue tropism.

The approach we took in this study focused on identifying a molecular feature that was shared across K-Ras non-permissive tissues. We find that low baseline expression of c-Myc may be this common denominator. We do not find that the degree of MAPK activation predicts permissivity, arguing against the general applicability of the ‘sweet spot’ model^55^ in this particular setting. Instead, we propose that in the absence of c-Myc, K-Ras signaling is cut short downstream of ERK phosphorylation, resulting in the absence of a pro-proliferative response and a lack of a Ras overactivation phenotype in non-permissive tissues. Consequentially, we show that ectopic expression of c-Myc is sufficient for mutant K-Ras to elicit an oncogenic phenotype in the liver. Nevertheless, factors downstream of c-Myc may act as additional gatekeepers in other tissue contexts. For example, ectopic expression of c-Myc is unable to promote proliferation in the adult heart because Cyclin T1, a critical mediator of this response, is not expressed^56^. We hypothesize that a similar mechanism is at play in the pancreas, which is non-permissive to K-Ras^G12D^ despite moderate c-Myc expression levels. In this tissue, inflammation-induced epigenetic changes in the signaling network downstream of c-Myc may be required for oncogenic K-Ras signaling to be propagated.

Our results also indicate that additional, c-Myc independent, resistance mechanisms to oncogenic K-Ras exist. As discussed above, p53-dependent cell cycle arrest in response to K-Ras^G12D^ expression appears unique to skeletal muscle; and occurs despite an only moderate increase in ERK activation. Whether and how Ras signals in this tissue independently of c-Myc in a p53-mutant setting remains to be explored. Furthermore, low expression of Ras itself may be a limiting factor of oncogenic signaling in the adult heart. Altogether, it is possible that no unifying mechanism of tissue permissivity to oncogenic K-Ras – or indeed any other cancer driver – exists. In other words, not all non-permissive tissues may be non-permissive for the same reasons and the same is true for permissive tissues.

Our understanding of the interplay between oncogenic K-Ras and endogenous c-Myc is limited. Although c-Myc and K-Ras are often cited as a prominent example of oncogene cooperativity, much of this interdependence is gleaned from experiments using ectopic expression of mutant Ras; a context that requires c-Myc overexpression to inhibit Ras-induced cell cycle arrest to allow for cellular transformation^57^ and a scenario that is not recapitulated in the setting of oncogenic K-Ras expression at endogenous levels^47^. The investigation of Myc and K-Ras cooperativity *in vivo* has primarily focused on the contribution of augmented c-Myc expression to the promotion of K-Ras mutant tumor progression in the lung and pancreas^41,58,59^. More pertinent in the context of K-Ras permissivitity in the absence of Myc amplification, however, are studies that demonstrated that endogenous c-Myc is essential for the development and maintenance of K-Ras-driven lung adenomas^60,61^. Analogously, we show that the molecular network of the liver is capable of driving K-Ras oncogenicity, but only when c-Myc – a component that is missing from the network at steady state – is introduced into the system. Collectively, the data presented here support the notion that c-Myc is an obligate node of oncogenic K-Ras signaling in all tissues, and that its baseline expression levels principally shape the tropism of mutant K-Ras.

On a general level, our results highlight that oncogenes cannot exert pro-tumorigenic effects in isolation, but instead tap into the pre-existing signaling network of a cell. By extension, the tissue-specific effects of oncogene expression are necessarily determined by the presence or absence of essential downstream effectors.

## SUPPLEMENTAL INFORMATION

Document S1. Figures S1-S7

Table S1. Ten tissue shotgun proteomics data set, related to Figure 2.

Table S2. Ten tissue RAS pathway targeted proteomics data set, related to Figure 4

## ACKNOWLEDGEMENTS

This work was supported by an award from the Cancer Research UK Grand Challenge and the Mark Foundation to the SPECIFICANCER team. Part of this work was funded by the National Institutes of Health grant CA282268 (Q.Y.), and the NIH/NCI P30 grant CA008748 (C.S.). K.M.H. is also supported by the Dana-Farber Cancer Institute Hale Center for Pancreatic Cancer Research and the Project P fund.

The authors would like to thank the Neurobiology Imaging Facility at Harvard Medical School, Boston, MA for Imaging support. Portions of this research were conducted on the O2 High Performance Compute Cluster, supported by the Research Computing Group, at Harvard Medical School.

For their technical assistance with various parts of this study we would like to thank Caroline Fahey, Ariel Karson, Sarah Schwarz, Deborah Burkhart, Beatrice Awasthi, Joshua Cook, Utsarga Adhikary, and Salva Casaní-Galdón.

## AUTHOR CONTRIBUTIONS

Conceptualization: O.P. and K.M.H.; Methodology: O.P., Q.Y., C.Y., Y.K., K.M.H.; Software: C.Y.; Formal analysis: O.P., Q.Y., C.S.; Investigation: O.P., Q.Y., S.H., J.A.P., B.H., S.X.; Resources: C.P.P., S.P.G., K.M.H.; Writing – Original Draft: O.P.; Writing – Review & Editing: O.P. and K.M.H.; Visualization: O.P.; Supervision: O.P., Y.K., K.M.H.; Funding acquisition: K.M.H.

## METHODS

### EXPERIMENTAL MODEL AND STUDY PARTICIPANT DETAILS

#### Animal Models

Wild-type C57BL/6J (IMSR_JAX:000664), *ROSA26*^Cre-ERT2^ (IMSR_JAX:008463)^12^, UBC-CreERT2^TG^ (IMSR_JAX:007001)^13^, *ROSA26*^LSL-EYFP^ (IMSR_JAX:006148)^14^, *Myh6*-MerCreMer^TG^ (IMSR_JAX:005657)^15^, and *ROSA26*^CAG-LSL-MYC^ (IMSR_JAX:0020458)^40^ mice were obtained from Jackson Laboratory. *Kras*^LSL-G12D^ (IMSR_NCIMR:01XJ6)^11^ mice were obtained from the NCI Mouse Repository. *Kras*^LSL-WT^ mice were generated by blastocyst microinjections performed by the Transgenic Mouse Core at Harvard Medical School using *Kras*^LSL-WT^ ES cells^62^ that were a gift from David Tuveson and a byproduct from the generation of *Kras*^LSL-G12D^ mice. The LSL-WT locus in these mice is identical to the LSL-G12D locus in *Kras*^LSL-G12D^ mice except that it is missing the single nucleotide substitution that encodes the G→D mutation. All mouse strains were maintained on a C57BL/6J background. Animals were house in a specific-pathogen-free facility with 12-hour light-dark cycles and food and water available *ad libitum*. Female and male animals – used in largely equal proportions – were enrolled in experiments between 8-12 weeks-of-age. All animal work was reviewed and approved by the Institutional Animal Care and Use Committee (IACUC) at Beth Isreal Deaconess Medical Center or Dana Farber Cancer Institute. Animals were cared for, and experiments were performed according to IACUC guidelines.

## METHOD DETAILS

### Animal treatments

#### Tamoxifen

Tamoxifen (MilliporeSigma, #T5648) was first dissolved in 100% ethanol to 50 mg/mL and then further diluted in corn oil to a final concentration of 10 mg/mL in 20% ethanol/80% corn oil. Aliquots were stored at −20°C and thawed to 37°C immediately prior to use. For recombination efficiency analysis (Figure S1A) and short-term experiments, animals were injected intraperitoneally with 1 mg (female) or 1.3 mg tamoxifen (male) once a day for six consecutive days to induce Cre-mediated recombination in as many target cells as possible. Because the overall body condition of UBC-CreER;R26-YFP;*Kras*^LSL-G12D^ mice undergoing this treatment deteriorated quickly, tissues from all animals of this experimental cohort were harvested seven days after the last tamoxifen injection. For long-term experiments animals were injected once intraperitoneally with 1 mg (female) or 1.3 mg tamoxifen (male) to induce Cre-mediated recombination in a subset of target cells. Tissues from UBC-CreER;R26-YFP;*Kras*^LSL-G12D^ undergoing this treatment regimen were harvested when animals reached a human endpoint, which occurred 17-33 days after tamoxifen injection. Tissues from control *Kras*^LSL-WT^ mice were harvested 22-30 days after tamoxifen treatment. Tamoxifen dosage was adjusted based on the approximately 1.3-times higher weight of male compared to female mice at 8-12 weeks-of-age^63^.

#### AAV8.Cre

Ready-to-use AAV8.TBG.PI.Cre.rBG (AAV8.Cre) particle preparations were purchased from Addgene (#107787-AAV8) and diluted with sterile PBS immediately before use. Animals were injected once intravenously via the tail vein with 100 μl AAV8.Cre/PBS. To induce recombination in ∼100% of hepatocytes (see ^64^), mice were injected with 2×10^11^ genetic copies (GCs) of AAV8.Cre. These mice were harvested after 13 days (short-term). To induce recombination in ∼10% of hepatocytes (determined experimentally using ddPCR), male mice were injected with 0.5×10^10^ GCs and female mice with 1×10^10^ GCs of AAV8.Cre.

#### EdU

5-etyhyl-2’-deoxyuridine (EdU, Thermo Fisher Scientific, #A10044) was dissolved to 2.5 mg/mL in sterile PBS. Aliquots were stored at −20°C and thawed to 37°C immediately prior to use. One hour prior to tissue harvest, mice were injected intraperitoneally with 10 μl of diluted EdU per g body weight.

#### CCl_4_

Carbon tetrachloride (CCl_4_, MilliporeSigma, #319961) was diluted to 20% in corn oil immediately before use. Mice were injected once intraperitoneally with 2.5 μl CCl_4_/corn oil per g body weight.

#### EGF and PD0325901 (Mirdametinib)

Recombinant mouse EGF (Fisher Scientific, #2028EG200) was dissolved to 0.4 mg/mL in sterile PBS and administered by retro-orbital injection 5 min prior to tissue harvest under isoflurane anesthesia at 1 mg/kg. The MEK inhibitor PD0325901 (Selleck, #S1036) was dissolved to 90 mg/mL in DMSO/0.5% hyroxypropylmethyl cellulose and administered to mice one hour prior to tissue harvest by oral gavage at 10 mg/kg.

### Tissue harvest and processing

For MS-based protein analysis and baseline c-Myc protein quantification in tissues from wild-type C57BL/6J mice (8 weeks-of-age, purchased from The Jackson Laboratory), animals were euthanized by cervical dislocation and all tissues were harvested and snap frozen in liquid nitrogen within four minutes. Colons and small intestines were flushed with PBS prior to freezing. Eight-week old wild-type mice treated with EGF or PD0325901 (one mouse per treatment) were also euthanized by cervical dislocation prior to tissue harvest. For all other downstream analyses mice were euthanized by CO_2_ overdose followed by cervical dislocation. For staining preparations tissues were fixed in 3.7% formaldehyde (MilliporeSigma, #F1635) in PBS for 24 hours at RT and then transferred into 70% ethanol prior to standard paraffin embedding (performed by the Beth Israel Deaconess Medical Center Histology Core, Boston, USA). Tissues from *ROSA26*^Cre-ERT2*/LSL-EYFP*^, UBC-CreERT2^TG^;*ROSA26^LSL-EYFP/+^*, and *Myh6*-MerCreMer^TG^;*ROSA26^LSL-EYFP/+^*mice used for recombination analysis and tissues from EGF/PD0325901-treated mice for the generation of anti-phosphorylated ERK1/2 staining controls were harvested for fixation and downstream analysis by IF staining only. Tissues from all other mice were divided and one part of the organ was fixed in formaldehyde, and one part designated for protein/DNA/RNA extraction was snap frozen in liquid nitrogen. Colons and small intestines were flushed with PBS, placed on filter paper and opened longitudinally. When applicable, longitudinal sections from colon and small intestine were removed and frozen, and the rest was fixed flat with the luminal side facing upward. Lungs were inflated with 3.7% formaldehyde/PBS, if part of the organ needed to be reserved for protein/DNA/RNA analysis the right lobes were clamped off using hemostatic forceps before being removed and frozen.

Frozen tissues were crushed on dry ice using a porcelain pestle and mortar (Coorstek, #60311). A maximum of 0.5 mL of sample was transferred to 2 mL safe-lock tubes together with four pre-cooled 05 mm stainless steel grinding balls (Retsch, #05.368.0034). Samples were pulverized in a cryogenic mixer mill (CryoMill, Retsch, #20.749.0001) equipped with a 2 mL reaction vial adapter (Retsch, #02.706.0303) using three (or two, for sample volumes <0.1 mL) 30 s grinding cycles at 20 Hz alternating with 30 s intermediate cooling cycles at 5 Hz. The first grinding cycle was preceded by a 2 min pre-cooling cycle at 5 Hz. Before placing the first set of samples into the mill, the cooling jacket and grinding jar were pre-cooled using the automatic pre-cooling cycle.

### Protein analysis

#### ELISA

Powdered tissue was resuspended in ice-cold 1× FastScan™ ELISA Cell Extraction Buffer (Cell Signaling Technology (CST), #69905) supplemented with 1× Enhancer Solution (CST, #25243), 1× protease/phosphatase inhibitor cocktail (CST, #5872), and 1× cOmplete™ EDTA-free protease inhibitor cocktail (MilliporeSigma, #11836170001). Lysates were rotated end-over-end for 30 min at 4°C, passed ten times through a syringe with a 22G needle (on ice), and centrifuged at 14,000×g for 30 min at 4°C. Supernatants were frozen in liquid nitrogen and stored at −80°C until further processing. Protein concentrations were determined using the Pierce bicinchoninic acid (BCA) Protein Assay Kit (Thermo Fisher Scientific, #23227). FastScan™ ELISA kits were used to detect total ERK1/2 (CST, #67404) and phosphorylated ERK1/2 (CST, #42173) following manufacturer’s instructions. 15 μg of protein/well was used for total ERK1/2 detection for all tissues except brain. To avoid assay saturation due to high expression levels of ERK1/2 in the brain, 4.8 μg of protein/well was used for this tissue. For phospho-ERK1/2 detection, 60 μg of protein/well was used for all tissues. TBP substrate incubation was performed at RT with agitation. Absorbance at 450 nm was read using the GloMax® Explorer Microplate Reader (Promega, #GM3500). Measurements were performed in technical duplicates or triplicates and averaged.

#### Ras RBD pull-down assays

GST-tagged Ras-binding domain of C-Raf (Raf-RBD; a gift from Channing Der, Addgene_13338) was expressed in BL21-AI cells as described by Taylor et al.^65^ with changes adapted from Johnson et al.^66^ After induction with 1mM Isopropyl β-D-1-thiogalactopyranoside (IPTG) bacteria were grown at 37°C for 4 h with shaking. Bacterial pellets were washed once in HBS buffer, pH 7.5 (25 mM HEPES, 150 mM NaCl) and resuspended in RLB buffer, pH 7.5 (20 mM HEPES, 120 mM NaCl, 10% glycerol) supplemented with 1× cOmplete™ EDTA-free protease inhibitor cocktail (MilliporeSigma, #11836170001). Bacteria were lysed using a microfluidizer (Microfluidics, #M110-L). Lysates were cleared by centrifugation at 10,000 rpm for 10 min at 4°C. Aliquots were frozen in liquid nitrogen and stored at −80°C until further use.

Powdered tissue was resuspended in ice-cold MLB buffer, pH 7.5 (25 mM HEPES, 150 mM NaCl, 1% IGEPAL® CA-630 (MilliporeSigma, #18896), 0.25% sodium deoxycholate, 10% glycerol, 10 mM anhydrous MgCl_2_) supplemented with 2× cOmplete™ EDTA-free protease inhibitor cocktail, 1% phosphatase inhibitor cocktail 2 (MilliporeSigma, #P5726), and 1% phosphatase inhibitor cocktail 3 (MilliporeSigma, #P0044). Lysates were rotated end-over-end for 30 min at 4°C and cleared by centrifugation at 14,000×g for 30 min at 4°C. Lysates were frozen in liquid nitrogen and stored at −80°C. Tissue lysates were thawed on ice immediately before use in Ras RBD pull-downs.

For binding of Raf-RBD to glutathione beads, bacterial lysates were thawed on ice. IGEPAL® CA-630 was added to a final concentration of 0.5% and mixed by inversion. Glutathione Sepharose™ resin (Cytiva, #17075601) was washed twice with RLB buffer and then incubated with Raf-RBD bacterial lysate for one hour with end-over-end rotation at 4°C. Raf-RBD-bound resin was washed five times with RLB buffer containing 0.5% IGEPAL® CA-630, distributed between 1.5 mL reaction tubes for Ras RBD pull-downs (∼20 μl beads/tube), and kept on ice. Bacterial lysates from a single preparation were used for all Ras RBD pull-down assays in this study. Raf-RBD concentration was estimated by Coomassie staining.

Tissue lysates were incubated with Raf-RBD-bound resin in 500 μl MLB buffer supplemented with 1× cOmplete™ EDTA-free protease inhibitor cocktail for one hour with end-over-end rotation at 4°C. One mg of protein input was used per pull-down (protein concentrations were determined by BCA assay). After incubation with tissue lysate, resin was washed three times with ice-cold MLB buffer. The last wash was aspirated, leaving ∼30 μl buffer behind. Ten μl of NuPAGE™ LDS Sample Buffer (Thermo Fisher Scientific, #NP0007) was added and samples were frozen in liquid nitrogen and stored at −80°C until analysis by SDS-PAGE.

#### TCA precipiation

For the detection of (phospho-)c-Myc in tissue samples, protein was precipitated using trichloroacetic acid (TCA) to prevent degradation (inspired by Spiller and Tidd^39^). Powdered tissue (<3 mm^3^) was added to 1 mL ice-cold 20% TCA, vortexed, and incubated on ice for one hour. Precipitated protein was pelleted by centrifugation at 14,000 rpm for 20 min. Protein pellets were washed twice with 100% acetone, and once with 100% methanol, both chilled to −20°C. All centrifugation steps were performed at 4°C. After the last wash, protein pellets were air dried for 5 min. Ninety μl of RIPA buffer (Boston BioProducts, #BP-115) supplemented with 1× cOmplete™ EDTA-free protease inhibitor cocktail, 1% phosphatase inhibitor cocktail 2 (MilliporeSigma, #P5726), and 1% phosphatase inhibitor cocktail 3 (MilliporeSigma, #P0044) and 30 μl of 4× NuPAGE™ LDS Sample Buffer (Thermo Fisher Scientific, #NP0007) was added per pellet. Samples were heated to 95°C for 20 min, frozen in liquid nitrogen and stored at −80°C until analysis by SDS-PAGE.

#### SDS-PAGE and Western Blotting

Protein expression was generally measured in tissue lysates prepared in ELISA buffer (see above), except for Western Blots that included c-Myc detection (these samples were prepared by TCA precipitation). Protein lysates in ELISA buffer were mixed with NuPAGE™ LDS Sample Buffer (final concentration 1×; Thermo Fisher Scientific, #NP0007) and NuPAGE™ Sample Reducing Agent (final concentration 1×; Thermo Fisher Scientific, #NP0004). Fifty μg of protein was used per sample. Four μl of reducing agent was added to Ras RBD pull-downs already mixed with sample buffer (the entire volume was loaded onto gels). Samples were heated to 70°C for 10 min and separated on 1.0 mm 4-12% NuPAGE™ Bis-Tris Mini or Midi Protein gels (Thermo Fisher Scientific, #various) using NuPAGE™ MOPS SDS Running Buffer (Thermo Fisher Scientific, #NP000102) and NuPAGE™ Antioxidant (Thermo Fisher Scientific, #NP0005) following manufacturer’s instructions at 200 V for ∼50 min. Gels were washed in 20% ethanol prior to protein transfer. Transfers were performed using the Trans-Blot® Turbo™ RTA Midi 0.2 μm Nitrocellulose Transfer Kit (Bio-Rad, #1704271) and Trans-Blot® Turbo™ Transfer System (Bio-Rad, #1704150) according to manufacturer’s instructions, applying the Standard SD 30 min transfer program. Protein content in TCA precipitated samples was estimated by separating 18 μl of sample mixed with 2 μl reducing agent (after heating to 70°C for 10 min) by SDS-PAGE as described above. Nitrocellulose membranes were stained with Revert™ 700 Total Protein Stain (LICOR Biosciences, #926-11010) following manufacturer’s instructions. Membranes were imaged on an Odyssey® CLx Imager (LI-COR, #9140) and protein signal in the 700 nm channel of each lane was measured using Image Studio™ software (LI-COR, v5.2.5). TCA precipitated samples were then re-analyzed by SDS-PAGE as described above using adjusted sample volumes.

To ensure equal protein loading for Western Blots that contained samples from more than one tissue type, membranes were stained with Revert™ 700 Total Protein Stain, following manufacturer’s instructions prior to blocking. Membranes were generally not destained, unless Alexa Fluor™ 680 secondary antibodies (ABs) were used for downstream detection. Raf-RBD was also detected by Revert™ 700 Total Protein Stain; but only membrane sections containing Raf-RBD – not Ras – were stained.

Prior to incubation with primary ABs, membranes were blocked in Intercept® (TBS) Blocking Buffer (LICOR Biosciences, #927-60001) for one hour at RT with gentle agitation. Primary ABs were diluted in Intercept® T20 (TBS) Antibody Diluent (LICOR Biosciences, #927-65001) and incubated o/n at 4°C with gentle agitation. The following primary ABs were used (dilutions are indicated in brackets): anti-vinculin (1:1000; CST, #13901), anti-GFP/YFP (1:5000; Abcam, #ab13970), anti-Ras (G12D mutant specific) (1:1000; CST, #14429), anti-Ras (1:1000; MilliporeSigma, #05-516), anti-p21^Cip1/Waf^ (1:1000; Abcam, #ab188224), anti-ERK1/2 (1:2000; CST, #4696), anti-phospho-ERK1/2 (Thr202/Tyr204) (1:1000; CST, #4377), anti-c-Myc (1:1000; Abcam, #ab32072), anti-phospho-c-Myc (Ser62) (1:1000; CST, #13748). After incubation with primary ABs, membranes were washed three times for 10 min in 1×TBS/0.1%TWEEN® 20 (TBS-T) at RT with gentle agitation. Membranes were incubated with secondary ABs diluted 1:10,000 in Intercept® T20 (TBS) Antibody Diluent for 1 h at RT. The following secondary ABs were used: goat anti-rabbit Alexa Fluor™ 800 (Thermo Fisher Scientific, #A32735), goat anti-mouse Alexa Fluor™ 680 (Thermo Fisher Scientific, #A-21058), goat anti-chicken Alexa Fluor™ 680 (Thermo Fisher Scientific, #A32934). After incubation with secondary ABs, membranes were washed three times for 10 min in TBS-T at RT with gentle agitation and imaged on an Odyssey® CLx Imager. Western Blot bands were quantified using Image Studio™ software.

#### Mass spectrometry-based shotgun proteomics

Powdered tissue harvested from 12 wild-type 8-week-old C57BL/6J mice (six males, six females) was lysed in 8 M urea in 200 mM EPPS, pH 8.5 supplemented with 2× cOmplete™ EDTA-free protease inhibitor cocktail (MilliporeSigma, #11836170001). Samples were passed ten times through a syringe with a 22G needle and centrifuged at 14,000×g for 5 min. Supernatants were frozen in liquid nitrogen and stored at −80°C until further processing. Protein concentrations were determined by BCA assay (Thermo Fisher Scientific, #23227). To balance mouse-to-mouse variability, equal protein amounts from four mice (two males, two females) were pooled for each tissue sample and diluted to a final protein concentration of 1 mg/mL. A total of 30 pooled samples (three pools per tissue type, ten tissues) were processed. Disulfide bonds were reduced with 5 mM Tris Carboxy Ethyl Phosphene (TCEP; Thermo Fisher Scientific, #77720) for 15 min at RT and alkylated with 10 mM iodoacetamide (Thermo Fisher Scientific, #A39271) for 30 min at RT in the dark. Excess iodoacetamide was quenched by addition of dithiothreitol (DTT) to a final concentration of 5 mM and incubated for 15 min at RT. Two hundred μL of each pooled sample (∼200 μg protein) was subjected to methanol-chloroform precipitation in 2 mL sample tubes. Per tube were sequentially added 800 μL of methanol, 200 μL of chloroform, and 600 μL of water. Samples were vortexed after each addition. Samples were centrifuged at 14,000×g for 1 min at RT and top and bottom liquid layers were aspirated leaving the protein disk intact. Protein disks were washed once with methanol and pelleted by centrifugation. Protein pellets were resuspended in 200 μL of 200 mM EPPS, pH 8.5 before the addition of 4 μg of Lysyl Endopeptidase (Fisher Scientific, #NC9223464). Initial protein digestion was allowed to occur o/n at RT with constant agitation. More complete protein digestion was achieved through the addition of 4 μg trypsin (Fisher Scientific, #PI90059) per sample and incubation for six additional hours at 37°C with constant agitation. One hundred μL of each pooled sample (∼100 μg protein) was mixed with 30 μL anhydrous acetonitrile (ACN; Thermo Fisher Scientific, #51101) prior to labeling with 0.2 mg of TMT10plex™ isobaric label reagent (Thermo Fisher Scientific, #90406) for one hour at RT (the same label reagent was used for pooled samples of the same tissue type). For the generation of the bridge, 30 μL of each pooled sample was combined (final volume: 900 μL), mixed with 270 μL anhydrous ACN and labeled with 0.72 mg of TMT11-131C label reagent (Thermo Fisher Scientific, #A34807) for one hour at RT. Excess TMT reagent was quenched with hydroxylamine (final concentration: 0.3%; Thermo Fisher Scientific, #90115). Each of the three TMTplexes was comprised of one pooled sample per tissue type (channel 1-10) and the bridge (channel 11). One μL of each pooled sample (grouped by TMTplex) and bridge was combined and desalted by StageTip. Samples were analyzed on a Q Exactive™ Mass Spectrometer (Thermo Fisher Scientific) to obtain total sum intensities of each channel and calculate normalization factors. These were then used to combine the pooled samples (grouped by TMTplex) at a 1:1:1:1:1:1:1:1:1:1 ratio, while keeping the volume of bridge constant across all three TMTplexes. Combined samples were dried by vacuum centrifugation, resuspended in 1% formic acid (FA; Fisher Scientific, #PI85178) and desalted on a C18 Sep-Pak, 100 mg (Waters, #WAT023590). Desalted samples were again dried by vacuum centrifugation and resuspended in 5% ACN/10 mM ammonium bicarbonate, pH 8. Samples were fractionated using BPRP HPLC^67^ and an Agilent 1260 pump equipped with a degasser and a UV detector (set at 220 and 280 nm wavelength). Peptides were subjected to a 50-min linear gradient from 5% to 35% acetonitrile in 10 mM ammonium bicarbonate, pH 8 at a flow rate of 0.6 mL/min over an Agilent 300Extend C18 column (3.5 μm particles, 4.6 mm ID and 220 mm in length). The peptide mixture was fractionated into a total of 96 fractions, which were consolidated into 24 super-fractions^68^, and of which 12 non-adjacent super-fractions were analyzed. Samples were acidified with 1% formic acid and vacuum centrifuged to near dryness. Each super-fraction was desalted via StageTip, dried again via vacuum centrifugation, and reconstituted in 5% ACN/5% FA for LC-MS/MS processing.

Approximately 2 μg of each sample was analyzed on an Orbitrap Fusion Lumos instrument (Thermo Fisher Scientific) coupled to an Easy-nLC 1200 UHPLC (Proxeon). The 100 µm capillary column was packed with 35 cm of Accucore 150 resin (2.6 μm, 150Å; Thermo Fisher Scientific). Samples were fractionated over a ∼75 min gradient of 3 – 25% ACN in 0.125% FA at a flow rate of 500 nL/min. For data acquisition, the scan sequence began with an MS1 spectrum (Orbitrap analysis; resolution 120,000; mass range 400-1400 m/z; automatic gain control (AGC) target 2.0E5; maximum injection time 100 ms). Precursors for MS2/MS3 analysis were selected using a Top10 method. MS2 analysis consisted of collision-induced dissociation (quadrupole ion trap analysis with turbo scan rate; AGC 1.4e^4^; normalized collision energy (NCE) 35; maximum injection time 600 ms). MS3 precursors were fragmented by high energy collision-induced dissociation and analyzed using the Orbitrap (NCE 65; AGC 2e^5^; maximum injection time 150 ms, resolution was 50,000). Real-time search was enabled via Orbiter and used the mouse UniProt database for on-line searching^69,70^.

Spectra were converted to mzXML via MSconvert^71^. Database searching included all entries from the mouse UniProt reference database (downloaded: August 2021). The database was concatenated with one composed of all protein sequences for that database in the reversed order. Searches were performed using a 50-ppm precursor ion tolerance for total protein level profiling. The product ion tolerance was set to 0.03 Da. These wide mass tolerance windows were chosen to maximize sensitivity in conjunction with Comet searches and linear discriminant analysis^32,72^. TMT labels on lysine residues and peptide N-termini +229.163Da), as well as carbamidomethylation of cysteine residues (+57.021 Da) were set as static modifications; oxidation of methionine residues (+15.995 Da) was set as a variable modification. Peptide-spectrum matches (PSMs) were adjusted to a 1% false discovery rate (FDR)^73,74^. PSM filtering was performed using a linear discriminant analysis^32^ and then assembled further to a final protein-level FDR of 1%^73^. Proteins were quantified by summing reporter ion counts across all matching PSMs^75^. Reporter ion intensities were adjusted to correct for the isotopic impurities of the different TMT reagents according to manufacturer specifications. The signal-to-noise (S/N) measurements of peptides assigned to each protein were summed and values were normalized so that the sum signal of all proteins combined was equal across channels (normalization was performed separately for each TMTplex).

Data processing was performed in Perseus (v2.0.10.0). Data was filtered to remove reverse hits, non-mouse contaminants and proteins with a summed signal <100 across all channels of a TMTplex. Data was log_2_ transformed and protein measurements were normalized across TMTplexes using the geometric mean of the signal in the bridge channels. Principal component analysis (PCA) was performed in Perseus on a filtered data set without missing values across all TMTplexes. For downstream analysis, data was filtered for valid values in all channels of at least one TMTplex (Table S1).

Correlation analysis was performed with R (v4.4.2) in R studio (v2024.09.1) using the cor.test() function with default arguments and method = “spearman”. Pre-ranked gene set enrichment analysis (GSEAPreranked) and Leading Edge Analysis was performed using the GSEA desktop application (v4.3.3). GSEAPreranked was performed using the Mouse MSigDB hallmark gene set collection (v2024.1), the Mouse UniProt IDs chip annotation file (v2024.1), and 4000 permutations. Defaults were used for all other settings. For plotting purposes FDR values of gene sets with FDR q-val = 0 (E2F Targets, Interferon Gamma Response, G2M Checkpoint, Interferon Alpha Response, Oxidative Phosphorylation, Fatty Acid Metabolism, Adipogenesis), were replaced with the q-value of the gene set with the lowest reported FDR (Allograft Rejection, FDR q-val = 4.24×10^-5^). Multi-pathway plots were generated using the R packages genekitr (v1.2.8) and geneset (v0.2.7).

#### Targeted Mass Spectrometry by TOMAHAQ/Tomahto

The tissue lysates prepared for shot gun proteomics were also used for targeted MS analysis. 200 μg of protein per sample was process as described above for untargeted mass spectrometric analysis, except that tissue samples were not pooled. A total of 120 samples (6 male and 6 female mice, 10 tissues each) were processed. Each TMTplex was comprised of the ten tissue samples of an individual mouse (channel 1-10) and the bridge (channel 11), which was identical to the bridge sample used for shot gun proteomics experiments. Ratio checks were performed on an Orbitrap Eclipse™ Tribrid™ mass spectrometer. Samples of each TMTplex were combined at a 1:1 ratio across channels 1-10, again keeping the volume of the bridge constant across all twelve TMTplexes. A second ratio check was performed, and the obtained normalization factors were used to adjust S/N measurements of the endogenous peptides across channels of individual TMTplexes (see below). Because some trigger peptides contained oxidized methionine residues, endogenous peptide mixtures were oxidized by incubation with 0.1% H_2_O_2_ in 30% ACN for one hour at RT.

Trigger peptides were purchased from JPT as SpikeTides™ or Maxi SpikeTides™. Trigger peptides were solubilized in 25% ACN/1% FA and further diluted to ∼1.5 μM in 5%ACN/1% FA. Peptides were combined in equal proportions and desalted on a SOLA™ HRP SPE Cartridge (Thermo Fisher Scientific, #60109-001). Mass spectrometric analysis of the trigger peptide mixture guided serial adaptation of the mixture to adjust amounts of peptides with low intensity. Trigger peptides were labeled with Super Heavy TMT label reagent (shTMT; Thermo Fisher Scientific, #A43073) at a mass ratio of 5:1 (TMT:peptides) in 200 mM EPPS, pH 8.5 for one hour at RT. Labeling reaction was then quenched with hyrdroxylamine and desalted, as described above. ShTMT-labelled trigger peptide mixture was spiked into TMT1-11-labelled endogenous peptide samples and analyzed by Triggered by Offset, Multiplexed, Accurate mass, High resolution, and Absolute Quantitation (TOMAHAQ)^29^. TOMAHAQ experiments were performed using the Tomahto software package^30,76^ on an Orbitrap Eclipse™ Tribrid™ mass spectrometer coupled to an Easy-nLC 1200 UHPLC system. Each sample was separated on an in-house packed C18 column (30 cm, 2.6 μm Accucore, 100 μm I.D.) and eluted using a 150-min method over a gradient from 5% to 38% B (95% acetonitrile, 0.125% formic acid). The instrument method only included Orbitrap MS1 scans (resolution at 120,000; mass range 300−1500 m/z; AGC target 2e^5^, max injection time 50 ms). Peptide targets were imported into Tomahto and the following decisions were made by Tomahto in real-time:

1. Tomahto listened to each collected MS1 scan.
2. When a precursor ion matched m/z of a potential trigger peptide (±10 ppm mass accuracy; matched charge state; minimal intensity of 5e^4^), Tomahto prompted insertion of an Orbitrap MS2 scan (<u>Trigger MS2</u>) with the trigger peptide’s precursor m/z (0.5 m/z isolation window; resolution at 15,000; AGC target 1e^4^; max injection time 120 ms; CID collision energy 35%). Once collected, a real-time peak matching strategy (RTPM) was used to confirm the identity of the trigger peptide (must match >6 fragment peaks within ±10 ppm).
3. If the trigger MS2 was successfully matched, Tomahto prompted the insertion of an Orbitrap MS2 scan (<u>Target MS2</u>) using the target peptide m/z (0.5 m/z isolation window; resolution at 15,000; AGC target 1e^5^; max injection time 900 ms; CID collision energy 35.1). The target peak m/z is a mixture of multiplexed endogenous peptides. At the same time, the MS2 fragment ions and their intensities for the trigger MS2 were stored in memory as a template library spectrum. After collection, the target MS2 scan was used to confirm that the target peptide was present at levels sufficient for detection. This was accomplished via RTPM where fragment ions must be present in the spectrum (±10 ppm) and rank ordered by intensity from the trigger MS2. SPS fragment ions were now selected from this scan. Only b- and y-type ions were considered for selection provided they had a TMT modification. SPS candidates were required to match the fragmentation pattern of the stored library spectrum, meaning fragment ratios relative to the highest fragment were within ±50% of that in the stored spectrum. In addition, each SPS candidate underwent a purity filter of 0.5 (at least 50% of the signal attributed to the fragment ion within a 3 m/z window) to be included in the final list.
4. Upon confirmation of target peptide presence and successful selection of SPS ions, Tomahto next triggered an ion trap SPS-MS3 prescan (normal scan mode; AGC target of 1e^6^; max injection time of 10 ms). This was used to quickly estimate the signal strength for the TMT reporter ions. This estimate was used to set the lengthy injection times needed for the SPS-MS3 scan detection in the Orbitrap.
5. Following the prescan, Tomahto prompted the insertion of the SPS-MS3 quantification scan (resolution of 50,000; SPS ions from part 2, 0.5 m/z window, max injection time of 5,000 ms).

Raw data were processed by the data analysis module of Tomahto. RawFileReader (Thermo Fisher Scientific) was used to read files and spectra were matched to synthetic trigger peptide or endogenous target peptides respectively. S/N intensities of endogenous target peptides corresponding to the same protein were summed and normalized across channels of individual TMTplexes using the normalization factor obtained from the secondary ratio check (see above). Downstream data processing was performed in Perseus (v2.0.10.0). Data was filtered to remove proteins with a summed signal <100 across all channels of a TMTplex. Data was log_2_ transformed and protein measurements were normalized across TMTplexes using the geometric mean of the signal in the bridge channels. Principal component analysis (PCA) was performed in Perseus on a filtered data set without missing values across all TMTplexes. For downstream analysis, data was filtered for valid values in all channels of at least one TMTplex (Table S2).

Correlation analysis was performed with R (v4.4.2) in R studio (v2024.09.1) using the cor.test() function with default arguments and method = “spearman”. Bonferroni correction was performed manually by multiplying p values by the number of performed tests (165). RAS pathway network shown in Figure 4 was manually generated in Cytoscape (v3.10.3) and modified in Adobe Illustrator.

For the detection of K-Ras4a (VEDAFYTLVR) and K-Ras4B (QGVDDAFYTLVR) peptides, trigger peptides were ordered from JPT as Maxi SpikeTides at >95% purity and labeled with shTMT reagent as described above. Trigger peptides were spiked into TMT1-11-labelled endogenous peptide samples (protein concentration 1 μg/μL) at a final concentration of 50 fmol/peptide. Samples were analyzed by TOMAHAQ/Tomahto and data was processed as described above. Spearman’s rank correlation coefficients were computed in Prism (v10.5.0).

### Droplet digital PCR

DNA was extracted from intact, crushed, or powdered frozen tissue (<5 mm^3^) using the *Quick*-DNA/RNA™ Miniprep Plus Kit (Zymo Research, #D7003). Tissue samples were added to 2 mL Lysing Matrix D tubes (MP Biomedicals, #116913500) containing 600 μL DNA/RNA Shield™ buffer and, if non-powdered tissue was used, homogenized for 20-30 min using a vortex with a tube adapter (Macherey-Nagel, #740469). Digestion with Proteinase K was performed for one hour at RT. All other steps were performed according to manufacturer’s instructions using 500 μL of cleared lysate. DNA concentrations were determined using a NanoDrop™ One Spectrophotometer (Thermo Fisher Scientific, #ND-ONE-W).

The primer and probe sequences for the detection of the recombined *Kras*^LoxP^ locus were adopted from Rakhit et al.^17^ and were identical except that the probe used here had 8 (instead of the original 9, used by Rakhit et al.) locked nucleic acids (LNAs). Droplet digital PCR (ddPCR) reactions were prepared with 1× ddPCR Supermix for Probes (Bio-Rad, #1863024), 1× Gt(ROSA)26Sor Copy Number FAM Assay (Bio-Rad, dMmuCNS592722274), 0.9 μM of each forward (5’-CCAGTCAACAAAGAATACCGCAAGG-3’) and reverse (5’-TCTGCATAGTACGCTATACCCTGTG-3’) *Kras*^LoxP^ primer, 0.2 μM HEX-labeled *Kras*^LoxP^ probe (5’-HEX-TCGACA<u>TA</u>A<u>C</u>T<u>TC</u>G<u>TAT</u>A-BHQ-1-3’, underlined bases denote LNAs; Sigma), and 15 ng of DNA in a reaction volume of 25 μL. Droplets were generated in the Automated Droplet Generator (Bio-Rad, #1864101) in 96-well PCR plates (Bio-Rad, #12001925) using droplet generation oil for probes (Bio-Rad, #1864110). Plates were sealed with foil (Bio-Rad, #1814040) using a PX1 PCR plate sealer (Bio-Rad, #1814000). PCR amplifications were performed on a C1000 Touch Thermal Cycler (Bio-Rad, #1851197) using the following conditions: 95°C for 10 min, 40 cycles of 94°C for 15 s and 57°C for 60 s, 98°C for 10 min, and 4°C hold. Droplets were analyzed on a QX200 Droplet Reader (Bio-Rad, #1864003) and data was processed with QX Manager Software, Standard Edition (v2.2).

*Kras*^LoxP^ fraction was calculated using the following equation: *Kras*^LoxP^ (copies per well) / 2× *Rosa26* CNV (copies per well). All samples were analyzed in technical duplicates and results were averaged.

### Tissue staining

Paraffin-embedded tissues were sectioned at five μm thickness, transferred onto microscope slides and dried for 12-24 hours at 40 °C. A portion of sections for this study were provided by the BIDMC Histology Core.

To enable the measurement of apoptotic cells (by TUNEL) and phosphorylated ERK1/2 localization (by IF) across multiple samples in a more cost-effective manner, we generated tissue microarrays (TMAs). These were constructed by the Brigham and Women’s Hospital (BWH) Histopathology Core using 3-4 cores (01.5-2 mm) per tissue block. Sections of TMAs were also provided by the BWH Histopathology Core.

Prior to staining, sections were deparaffinized for 2×10 min in Histo-Clear (National Diagnostics, #HS-202), followed by rehydration: 2×5 min 100% ethanol, 2×5 min 95% ethanol, 2×5 min 70% ethanol, and 2×5 min water.

#### Hematoxylin and Eosin (H&E) staining

Slides were stained for 45 s with Mayer’s Hematoxylin (Agilent Dako, #S330930-2), briefly rinsed in water, incubated in Bluing Reagent (Fisher Scientific, #6769001) for 1 min, washed in 70% and 95% ethanol (30 sec each), followed by staining in a freshly prepared 1:1 (v/v) Eosin (Fisher Scientific, #6766008):water solution for 22 seconds. Slides were dehydrated in 70% ethanol (2 min), 95% ethanol (2×2 min), 100% ethanol (5 min), and Histo-Clear (5 min). No. 1.5 coverslips were mounted to slides using Permount Mounting Medium (Fisher Scientific, #SP15-100). Slides were dried for a minimum of two days before bright-field images were acquired on a VS120-SL 5 slide scanner using a UPLSAPO 20× objective (Olympus). A portion of the H&E stainings for this study were provided by the BIDMC Histology Core using standard procedures.

Histopathologic evaluation of H&E-stained samples was performed by the Comparative Pathology Laboratory at the School of Veterinary Medicine, UC Davis. Liver samples from R26-MYC mice were evaluated by Carlie Sigel.

#### Immunofluorescence

Immunofluorescence stainings were performed following the protocol described by Robertson et al.^77^ with some modifications. De-paraffinized sections were immersed in 1× antigen retrieval solution, pH 9 (Agilent, #S236784-2) and heat-mediated antigen retrieval was performed for 30 min at high pressure in a pressure cooker. After manual pressure release, slides were allowed to cool within the staining well for one hour at RT. Slides were washed first in water (2×5 min) and then PBS (2×5 min). Sections were encircled with a hydrophobic marker (Fisher Scientific, #XT001PP) and transferred to a humidified chamber. For the generation of negative controls for anti-phosphorylated ERK1/2 staining, sections were incubated with 25.000 U/mL Lambda Protein Phosphatase in 1× NEBuffer for Protein MetalloPhosphatases and 1 mM MnCl_2_ (New England Biolabs, #P0753S) for one hour at RT before washing with PBS (3×5 min). Blocking was performed for one hour at RT in serum free blocking buffer (Agilent, #X090930-2). Primary antibodies were diluted in Antibody Diluent (Agilent, #S080983-2) and centrifuged for 5 min at 14.000 rpm before being transferred onto sections. The following primary ABs were used (dilutions are indicated in brackets): anti-GFP/YFP (1:250; Thermo Fisher Scientific, #A-11122), anti-CD45 (1:100; CST, #70257), anti-phospho-ERK1/2 (Thr202/Tyr204) (1:250; CST, #4370). Incubations with primary ABs were performed at 4 °C o/n. Sections were washed in PBS (3×5 min) prior to incubation with secondary AB for one hour at RT. Anti-rabbit secondary AB labeled with Alexa Fluor™ 647 (Thermo Fisher Scientific, #A-21244) was diluted 1:250 in Antibody Diluent and centrifuged for 5 min at 14.000 rpm before being transferred onto sections. Slides were washed once in PBS and incubated with 5 μg/mL (for spleen sections) or 1 μg/mL (for all other tissues) DAPI (BD Biosceinces, #P564907) for one hour at RT. DAPI dilutions were prepared in PBS and centrifuged for 5 min at 14.000 rpm before being transferred onto sections. After a final wash in PBS (3×5 min) no. 1.5 coverslips were mounted to slides using ProLong™ Diamond Antifade Mountant (Thermo Fisher Scientific, #P36970). Slides were dried for 24 hours at room temperature and transferred to 4 °C for short-term storage. Images were acquired on a VS120-SL 5 slide scanner (Olympus) equipped with a X-Cite® exacte fluorescence light source (Excelitas Technologies) and a Hamamatsu ORCA-R2 CCD camera (Spectra Services) using a UPlanSApo 20× objective (Olympus). All fluorescence microscopy for this study, including the imaging of EdU and TUNEL stainings (see below), was performed at the Neurobiology Imaging Facility at Harvard Medical School.

#### EdU staining

De-paraffinized sections were permeabilized by immersion in 0.5% Triton™ X-100 (MilliporeSigma, #X100) in PBS for 20 min at RT. Slides were washed twice in PBS. Sections were encircled with a hydrophobic marker and transferred to a humidified chamber. EdU detection was performed using the Click-iT™ EdU Cell Proliferation Kit for Imaging, Alexa Fluor™ 488 (Thermo Fisher Scientific, #C10337) following manufacturer’s instructions, except that all washing steps were performed with PBS. DNA staining with DAPI and mounting of cover slides was performed as described in the Immunofluorescence methods section. Images were acquired on a VS120-SL5 or VS200 slide scanner with a UPlanSApo 20× objective (Olympus).

#### TUNEL Assay

Terminal deoxynucleotidyl transferase-dUTP nick end labeling (TUNEL) was performed on de-paraffinized sections of TMAs using the Click-iT™ Plus TUNEL Assay, Alexa Fluor™ 488 (Thermo Fisher Scientific, #C10617) following manufacturer’s instructions. DNA staining with DAPI and mounting of cover slides was performed as described in the Immunofluorescence methods section. Images were acquired on a VS120-SL5 or VS200 slide scanner with a UPlanSApo 20× objective (Olympus).

### Image Analysis

Figure panels of microscopy images were generated in OMERO^78^. Quantification of markers detected by fluorescence microscopy was performed using a single nucleus segmentation analysis pipeline (see below), except for the quantification of EdU^+^ nuclei in the small intestine and colon, which was performed manually using OMERO.iviewer. Per mouse, a minimum of 50 crypts (long-term experiments) or 33-50 crypts (short-term experiments) located in the middle portion of the small intestine/colon were analyzed and the median number of EdU^+^ nuclei/crypt was calculated.

#### Single Nucleus Segmentation

Images were cropped to exclude non-relevant portions of tissues (e.g. pancreatic lymph nodes) and areas with excessive background fluorescence using Olympus VS-ASW (v2.9.2). Vsi files were stitched and converted into ome.tiff file format using NGFF-Converter (v1.1.4). Image analysis was performed using the MCMICRO pipeline^79^ on the O2 High Performance Compute Cluster at Harvard Medical School. Pre-stitched images from the Olympus slide scanner acquisition software were imported into MCMICRO where they were segmented first into probability maps (by UnMICST^80,81^ legacy) and nuclei label masks (by S3segmenter) using the DAPI channel as the nuclei marker. In UnMICST, the trained mouse model, ‘mousenucleiDAPI’, was used. In S3segmenter, nuclei were detected using a diameter range from 10 to 60 pixels. Individual parameter files are available upon request. Using the nuclei masks, mean fluorescence intensities of markers imaged in the second channel were measured and exported as single-nucleus data tables as .csv files. Fluorescence intensity thresholds were manually determined based on negative control slides that separated “positive” from “negative” nuclei in the second channel. Thresholds were applied to all images within one experimental batch (*i.e.* sections that were stained and imaged together). Percentages of “positive” nuclei were calculated separately for each section and averaged whenever stainings were performed in technical replicates (*i.e.* for slides with two serial sections). The number of nuclei analyzed per section varied between tissue types and was on average >600,000 (liver), ∼200,000 (brain), ∼270,000 (kidney), ∼160,000 (lung), ∼110,000 (pancreas), ∼36,000 (skeletal muscle), ∼360,000 (spleen), ∼49,000 (heart). For the quantification of apoptotic cells by TUNEL^+^ in TMAs, single-nucleus data were combined across cores from the same tissue sample. Per sample an average of 28,000 (liver), ∼34,000 (brain), ∼54,000 (kidney), ∼22,000 (lung), ∼19,000 (pancreas), ∼7,000 (skeletal muscle), ∼150,000 (spleen), ∼22,000 (heart), ∼21,000 (colon), ∼35,000 (small intestine) nuclei were analyzed.

## QUANTIFICATION AND STATISTICAL ANALYSIS

Statistical analysis was performed using Prism (v10.5.0), except for correlation analysis of shotgun and RAS pathway targeted proteomics data which was performed in R/RStudio as described in the Method Details. The type of statistical test performed and details regarding significance cut-offs are indicated in the figure legends associated with each experiment. All data points represent samples from individual mice, except for shotgun proteomics experiments for which samples from four individual mice were pooled (see Method Details). Box plots indicate 25^th^ to 75^th^ percentile and group median. Sample sizes were not predetermined through statistical methods.

## Supplemental Information for

**Figure S1.**
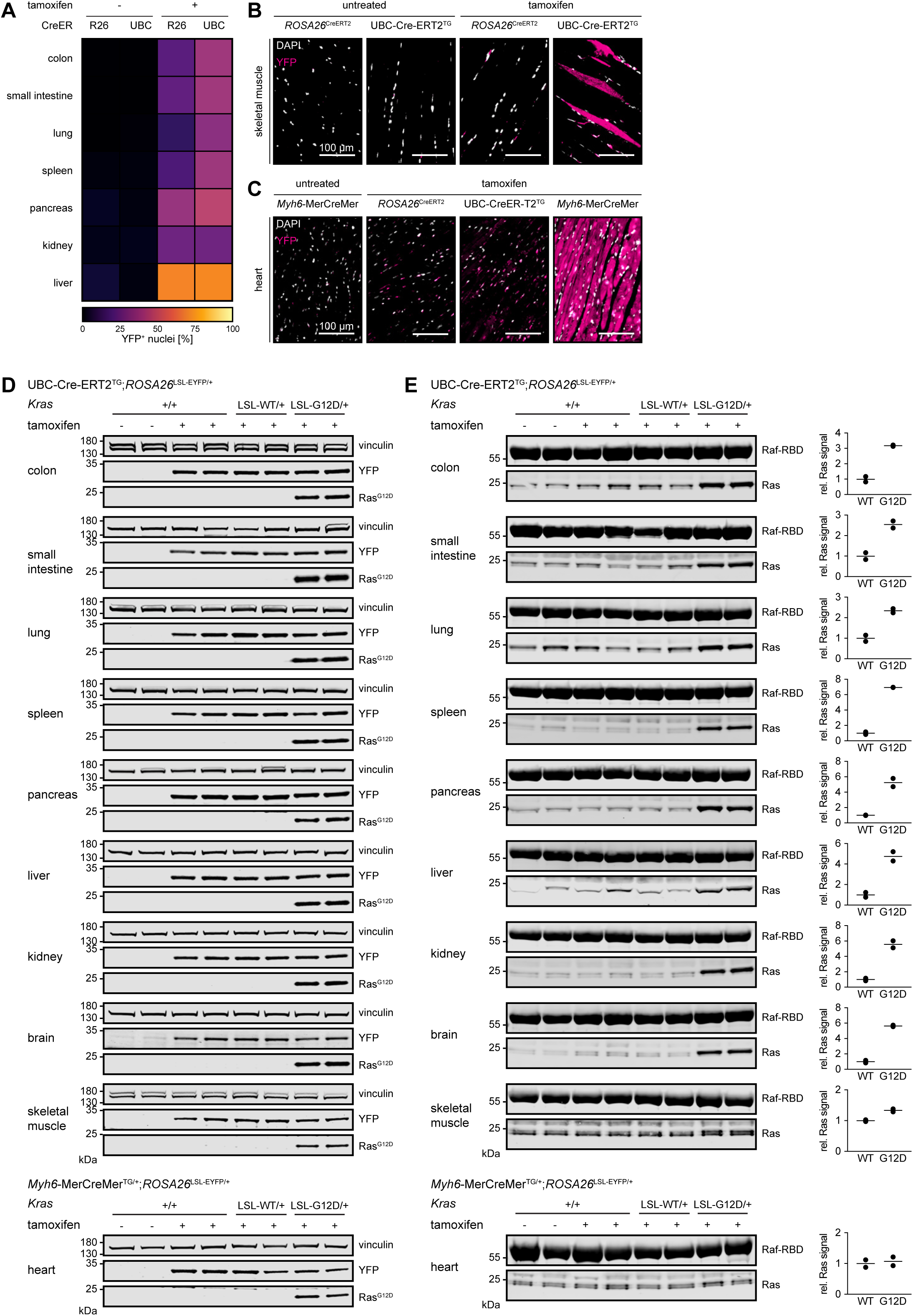
An experimental system for evaluating K-Ras tissue specificity, related to. Figure 1. (A) Recombination efficiency of the *ROSA26*^LSL-EYFP^ allele in mice expressing tamoxifen-inducible Cre recombinase from the *ROSA26* or UBC promoter. Tissues were harvested from untreated control animals and mice that received six consecutive doses of tamoxifen and that were harvested two weeks after. Cells in which recombination had occurred (YFP^+^) were quantified by single nucleus segmentation of whole slide immunofluorescence images. Shown is the median percentage of YFP^+^ nuclei across tissues from two male and two female mice. A considerable proportion of YFP^+^ nuclei were detected in the pancreata (4.48, IQR = 5.53, 2.80) and livers (7.05, IQR = 11.32, 6.77) of untreated R26-CreER/YFP mice indicating CreER leakiness in this model. Due to high background fluorescence, we were unable to accurately quantify YFP^+^ nuclei in the brain. (B and C) Because of non-specific nuclear fluorescence in a subset of skeletal and heart muscle cells, we assessed recombination efficiency in these tissues qualitatively. Shown are representative widefield fluorescence microscope images of skeletal (B) and heart (C) muscle from mice carrying the *ROSA26*^LSL-EYFP^ and the indicated Cre recombinase allele with and without tamoxifen treatment (six consecutive doses, harvested two weeks after) stained with DAPI (white) and anti-GFP/YFP (magenta). Scale bars = 100 μm. (D) Expression of YFP and Ras^G12D^ protein in tissues from mice with the indicated genotypes, untreated (-) or treated with six consecutive doses of tamoxifen (+) and harvested seven days after. (E) Levels of GTP-bound Ras evaluated by Raf-RBD pulldown. Same experimental cohort as in (D). Quantification of Ras signal in *Kras*^LSL-WT^ (WT) and *Kras*^LSL-G12D^ (G12D) samples is shown on the right.

**Figure S2.**
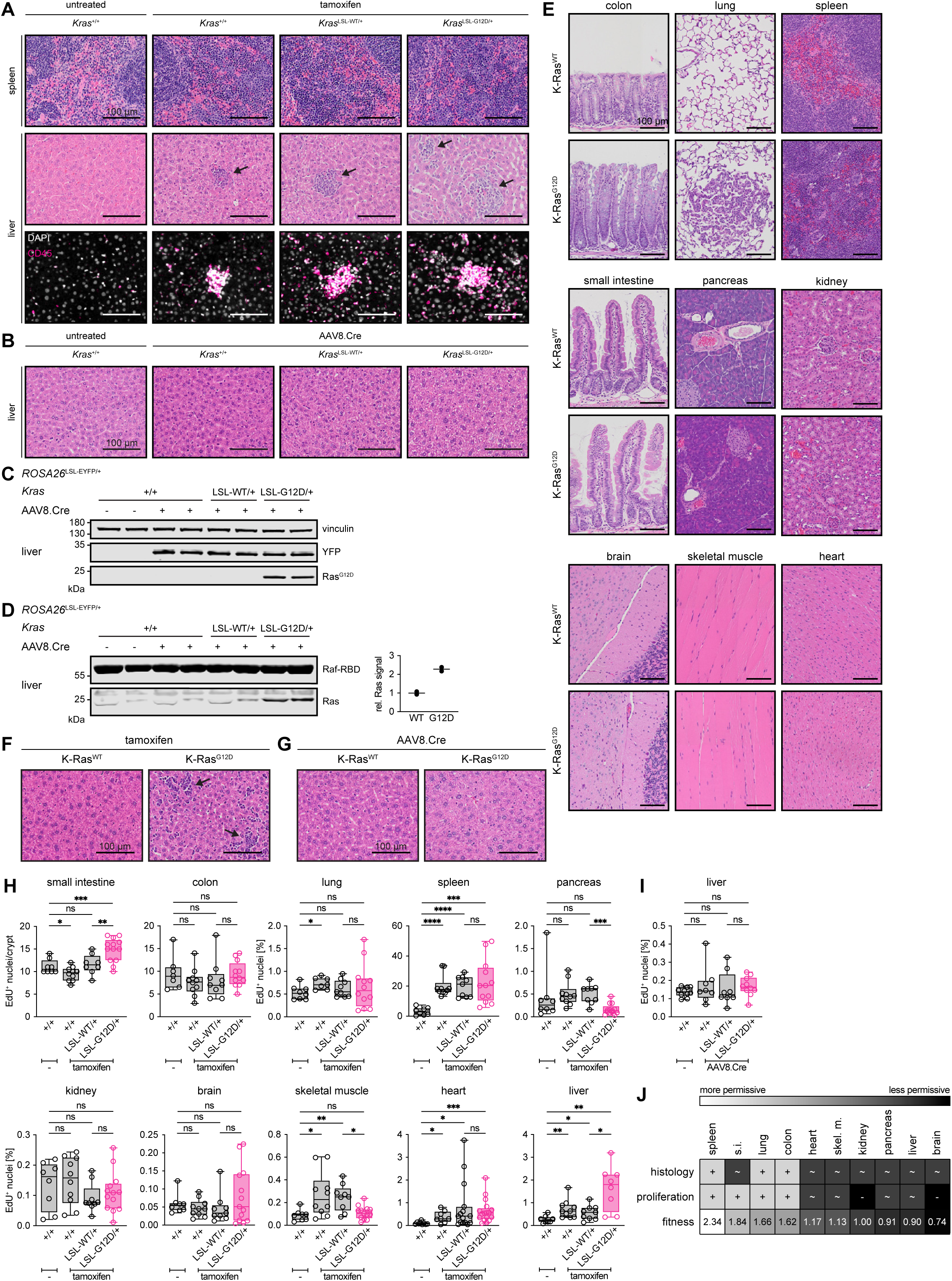
Expression of K-Ras^G12D^ confers an oncogenic phenotype only in a subset of tissues, related to Figure 1. (A) Representative images of H&E-stained sections of spleens (top) and livers (middle) from UBC-CreER;R26-YFP mice carrying the indicated *Kras* allele with and without tamoxifen treatment (six consecutive doses, harvested seven days after). Arrows indicate cellular infiltrates in the livers of tamoxifen-treated animals. Bottom panel: representative widefield fluorescence microscope images of livers from the same experimental cohort stained with DAPI (white) and anti-CD45 (magenta). Scale bars = 100 μm. (B) Representative images of H&E-stained sections of livers from R26-YFP mice carrying the indicated *Kras* allele injected with 2×10^11^ GCs of AAV8.Cre and untreated control animals harvested after 13 days. Note the absence of cellular infiltrates. Scale bars = 100 μm. (C) Expression of YFP and Ras^G12D^ protein in liver samples, same experimental cohort as in (B). (D) Levels of GTP-bound Ras evaluated by Raf-RBD pulldown in liver samples, same experimental cohort as in (B). Quantification of Ras signal in *Kras*^LSL-WT^ (WT) and *Kras*^LSL-G12D^ (G12D) samples is shown on the right. (E) Representative images of H&E-stained tissue sections from Myh6-CreER;R26-YFP (heart) and UBC-CreER;R26-YFP (all other tissues) mice carrying the conditional *Kras*-LSL-WT (K-Ras^WT^) or LSL-G12D (K-Ras^G12D^) allele treated with tamoxifen once and harvested after ∼28 days (long-term experimental cohort). Scale bars = 100 μm. (F) Representative images of H&E-stained sections of livers from UBC-CreER;R26-YFP mice, same experimental cohort as in (E). Arrows indicate cellular infiltrates. Scale bars = 100 μm. (G) Representative images of H&E-stained sections of livers from R26-YFP mice carrying the conditional *Kras*-LSL-WT (K-Ras^WT^) or LSL-G12D (K-Ras^G12D^) allele treated with AAV8.Cre (male: 0.5×10^10^ GCs, female: 1.0×10^10^ GCs) and harvested 28 days after. (H) Cell proliferation measured by EdU incorporation in tissues from Myh6-CreER;R26-YFP (heart) and UBC-CreER;R26-YFP (all other tissues) mice with (LSL-WT/+ or LSL-G12D/+) or without (+/+) a conditional *Kras* allele treated on six consecutive days with tamoxifen and harvested seven days after. Plotted is the median number of EdU^+^ nuclei per crypt (colon, small intestine; 33-50 crypts per mouse) or percentage of EdU^+^ nuclei (all other tissues). (I) Percentage of EdU^+^ nuclei in livers of R26-YFP mice with (LSL-WT/+ or LSL-G12D/+) or without (+/+) a conditional *Kras* allele treated with 2×10^11^ GCs of AAV8.Cre and harvested 13 days after. (H and I) Statistically significant differences between means were determined by Brown-Forsythe and Welch ANOVA tests. **** P < 0.0001, *** P < 0.001, ** P < 0.01, * P < 0.05, ns P > 0.05. (J) Summarized effects of K-Ras^G12D^ expression on histology, proliferation, and cell fitness across tissues. Presence of histological changes is denoted by “+”, absence by “∼”. Positive effect on proliferation is denoted by “+”, neutral effect by “∼“, and suppression by “-“. Mean fraction of *Kras*^LoxP-WT^ vs. *Kras*^LoxP-G12D^ cells at day 28 (see Figure 1E) was used to quantify the effect of K-Ras^G12D^ on cell fitness.

**Figure S3.**
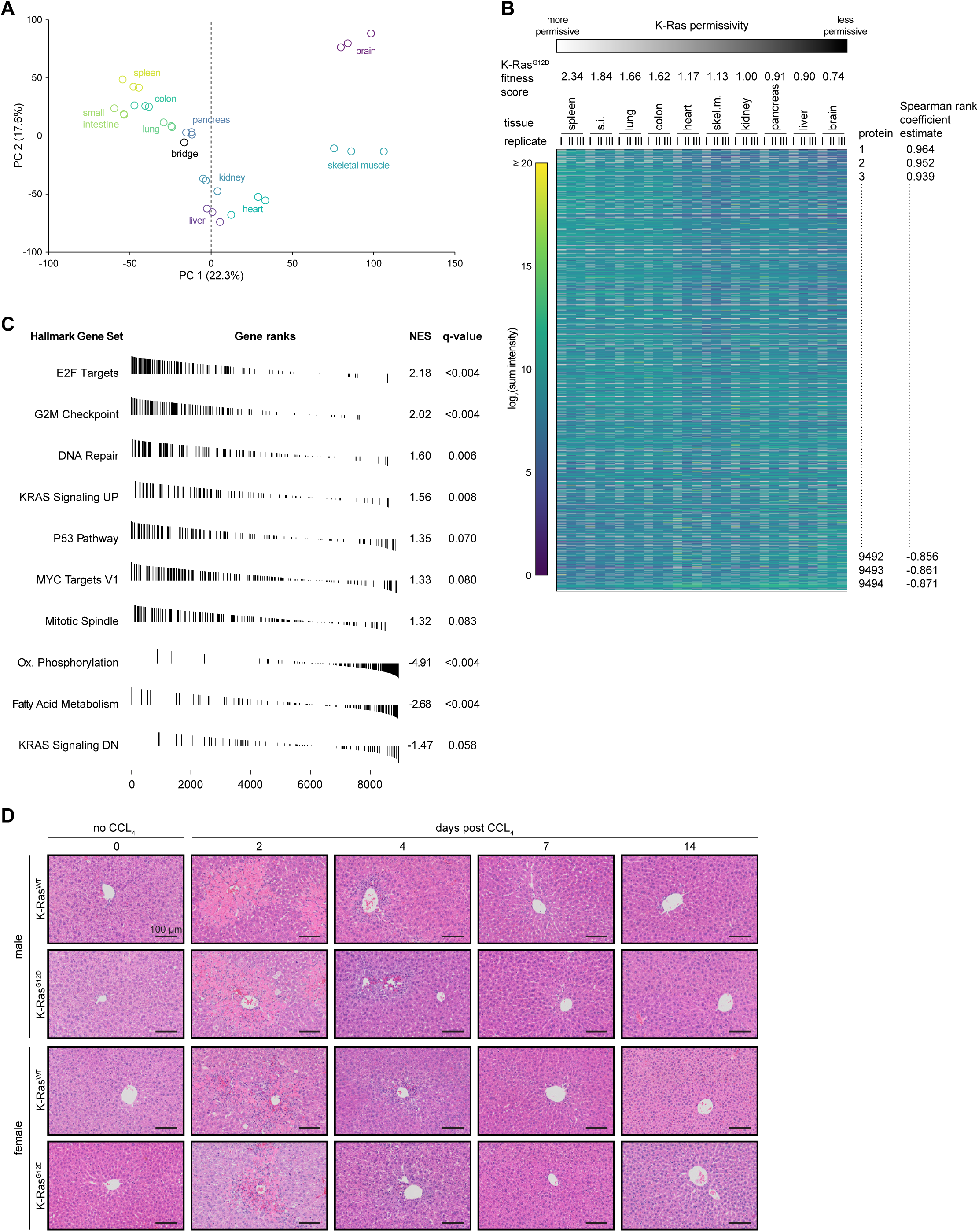
Interplay between cell proliferation and K-Ras permissivity, related to Figure 2. (A) Principal component analysis of global proteomics data set comprising ten tissues from wild-type mice across three TMTplexes. Plexes were normalized to a bridge channel, replicates of the bridge are thus projected on top of each other. (B) Tissues are ordered based on their K-Ras^G12D^ fitness score (see Figure S2J) from most (spleen) to least permissive (brain). Heatmap shows proteins ranked by the Spearman rank correlation coefficient between their relative expression (log_2_ TMT sum intensity) across tissues and the K-Ras permissivity of these tissues. (C) Gene ranks of select hallmark gene sets that were significantly (de-)enriched relative to tissue K-Ras permissivity by GSEAPreranked (see also Figure 2A). NES: normalized enrichment score, q-value: adjusted p-value controlling for false discovery rate (FDR). (D) Representative images of H&E-stained liver sections from R26-YFP mice carrying a *Kras*^LSL-WT^ (K-Ras^WT^) or *Kras*^LSL-G12D^ (K-Ras^G12D^) allele treated with 2×10^11^ GCs of AAV8.Cre at day −5, CCl_4_ at day 0 and harvested at indicated time points. Most left column shows liver histology in mice treated with AAV8.Cre but no CCL_4_. Centrilobular necrosis and lymphocytic infiltrates are evident at day two post CCL_4_ dosage with male mice (top two rows) generally displaying a more pronounced phenotype compared to female mice (bottom two rows). Tissue regeneration appears complete by day seven irrespective of *Kras* genotype. Scale bars = 100 μm.

**Figure S4.**
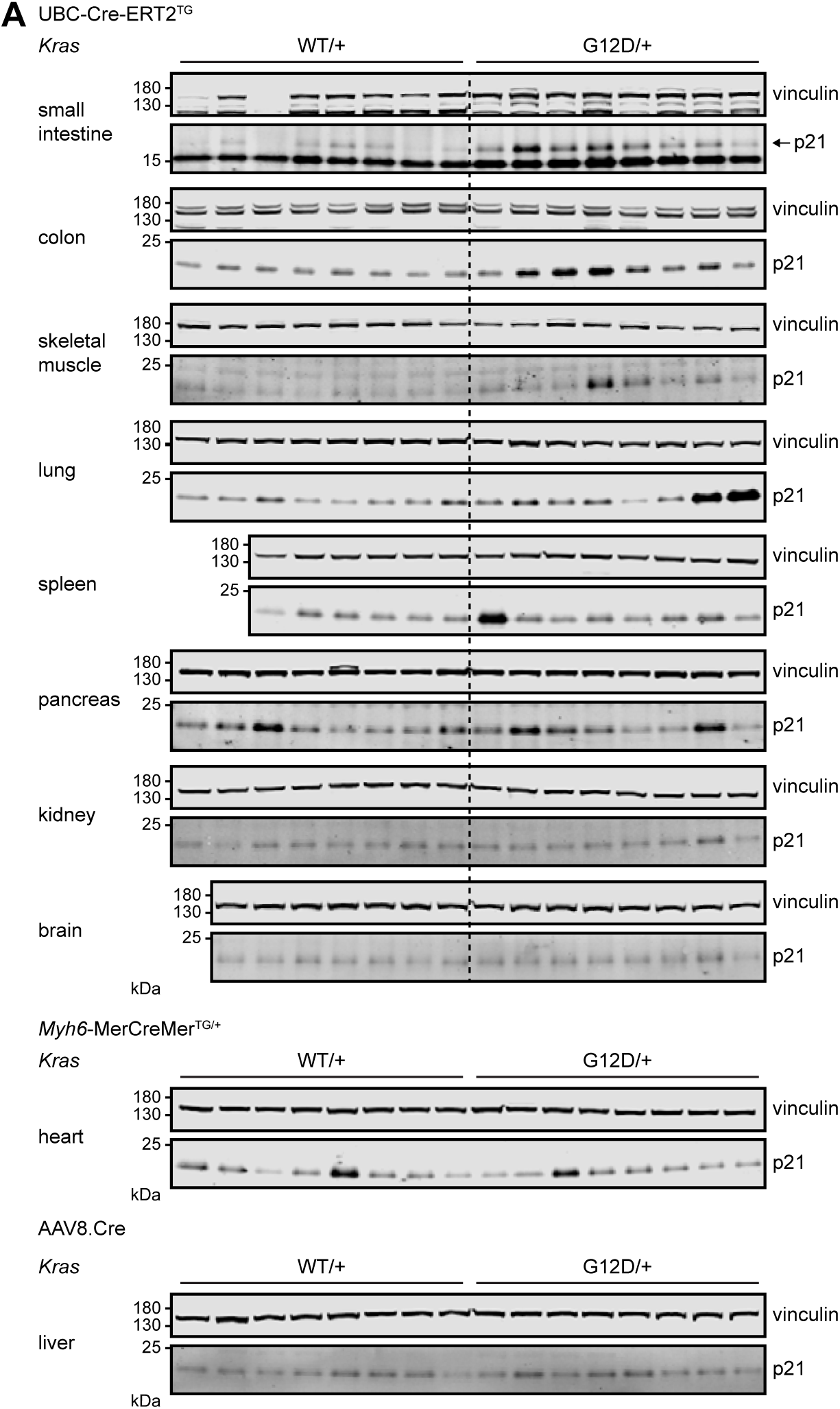
Effects of K-Ras^G12D^ expression on cell cycle arrest, related to Figure 3. (A) Assessment of p21^Cip1/Waf^ protein expression by Western blotting, quantification shown in Figure 3A. Tissue lysates are from R26-YFP (liver), Myh6-CreER;R26-YFP (heart), and UBC-CreER;R26-YFP (all other tissues) mice carrying a *Kras*^LSL-WT^ (WT/+) or *Kras*^LSL-G12D^ (G12D/+) allele injected with 2×10^11^ GCs of AAV8.Cre and harvested 13 days after (liver) or treated on six consecutive days with tamoxifen and harvested seven days after (all other tissues).

**Figure S5.**
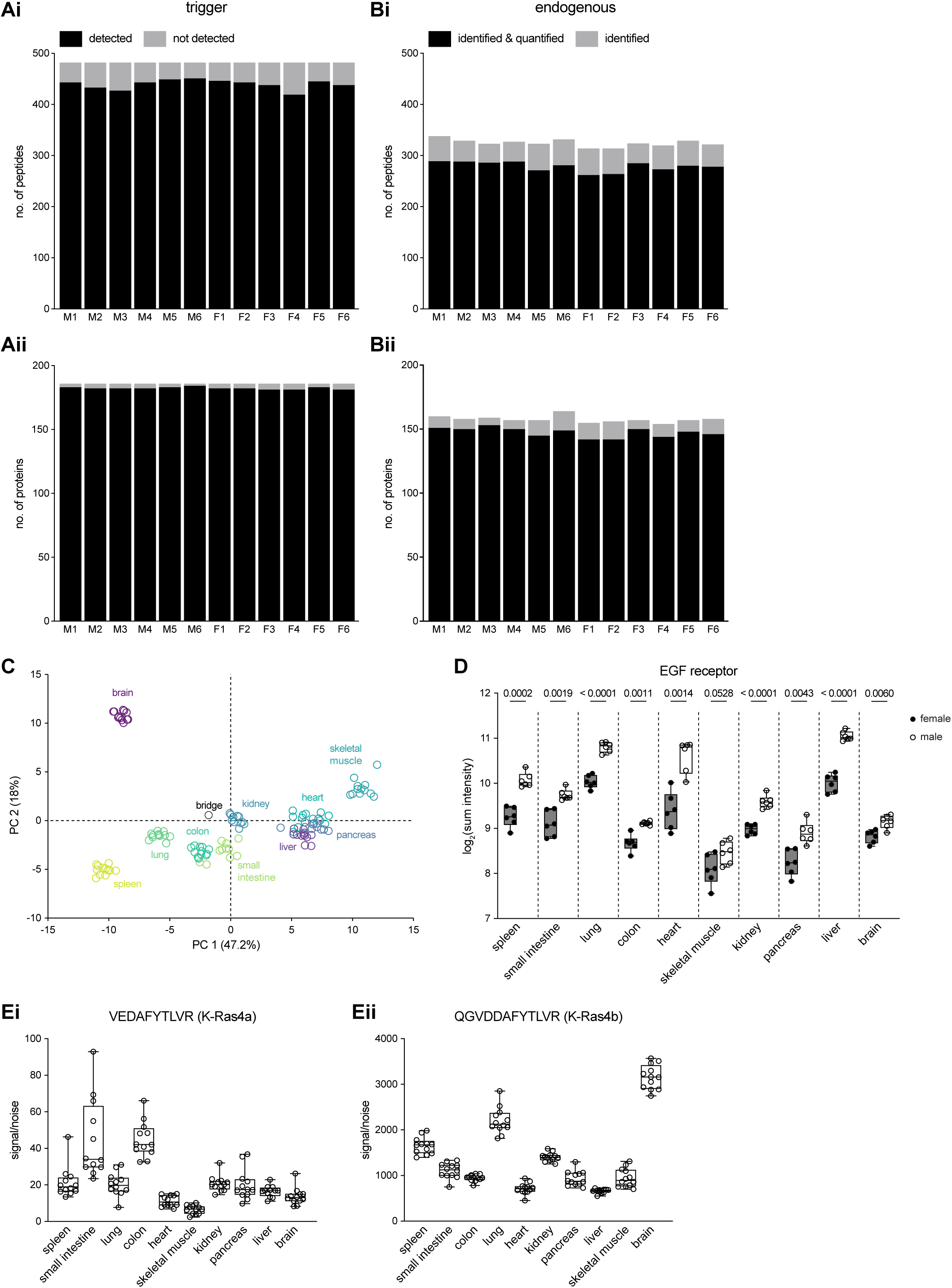
Relative quantification of members of the RAS pathway by targeted MS-based proteomics, related to Figure 4. (A) Number of detected (black) and missed (grey) trigger peptides (i) and corresponding proteins (ii) in each TMTplex. (B) Number of identified and quantified (black) and identified only (grey) endogenous peptides (i) and corresponding proteins (ii) in at least one channel in each TMTplex. (C) Principal component analysis of the targeted proteomics data set comprising ten tissues from wild-type mice across twelve TMTplexes. Plexes were normalized to a bridge channel, replicates of the bridge are thus projected on top of each other. (D) Relative protein expression of the epidermal growth factor receptor (EGFR) across wild-type tissues quantified by TOMAHAQ/Tomahto. Box plots indicate 25^th^ to 75^th^ percentile and group median. Differences in mean expression between female (filled circles) and male (empty circles) mice were evaluated with an unpaired t-test and the Holm-Šídák method for multiple comparison correction. P values are indicated above each comparison. (E) Relative signal-to-noise ratio of a K-Ras4a (i) and K-Ras4b (ii) specific peptide across wild-type tissues measured by TOMAHAQ/Tomahto. Each circle represents a measurement from an individual mouse/TMTplex. Box plots indicate 25^th^ to 75^th^ percentile and group median.

**Figure S6.**
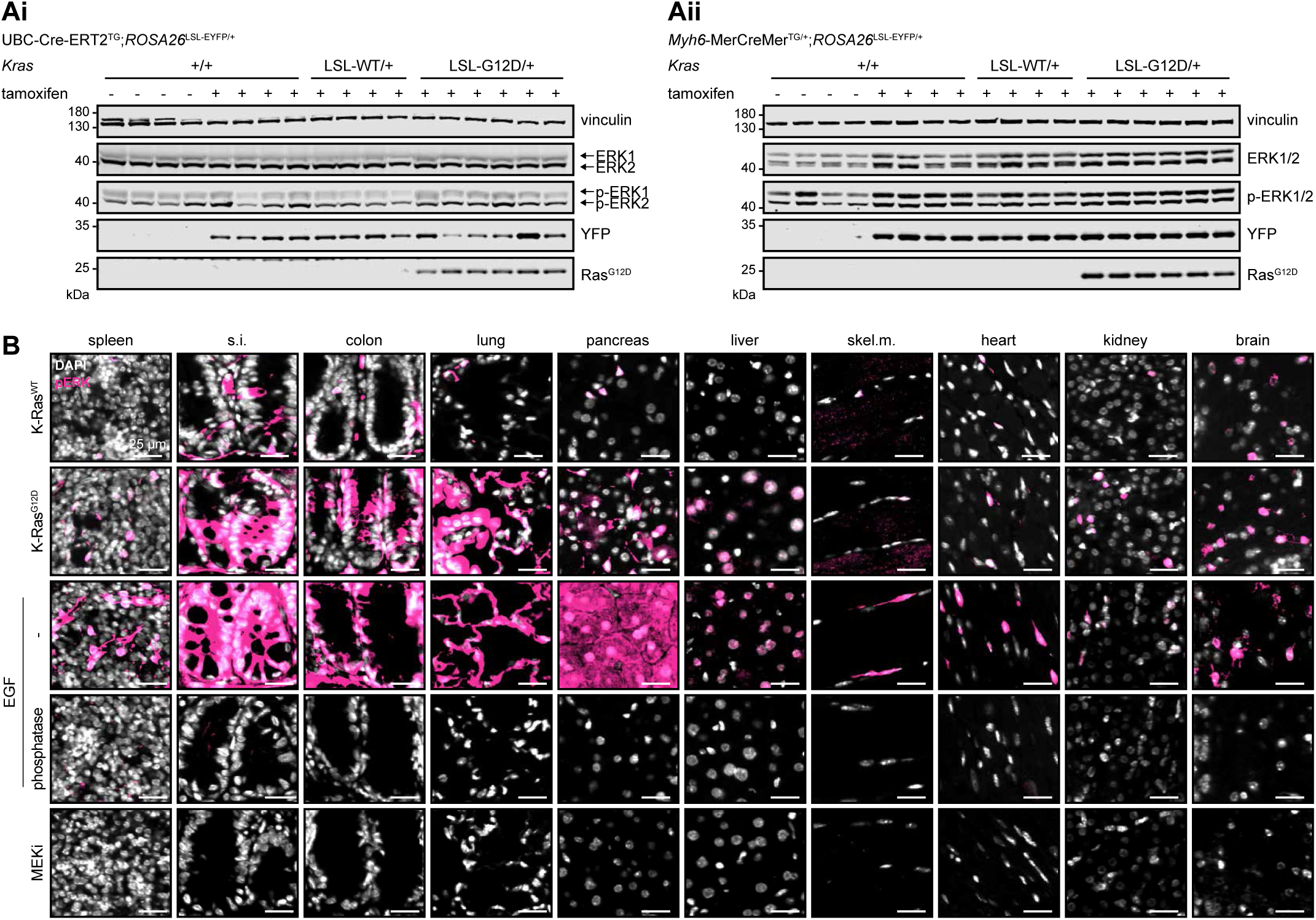
MAPK activation is not sufficient to mediate K-Ras permissivity, related to Figure 5. (A) Representative Western blot images used for the quantification of total and phosphorylated ERK1/2 protein expression in skeletal muscle (Ai) and heart (Aii) shown in Figure 5A. (B) Representative widefield fluorescence microscope images of tissue sections stained with DAPI (white) and anti-phospho-ERK1/2(Thr202/Tyr204) (pERK, magenta). Samples are from R26-YFP (liver), Myh6-CreER;R26-YFP (heart) and UBC-CreER;R26-YFP (all other tissues) mice carrying a *Kras*^LSL-WT^ (K-Ras^WT^) or *Kras*^LSL-G12D^ (K-Ras^G12D^) allele injected with 2×10^11^ GCs of AAV8.Cre and harvested 13 days after (liver) or treated on six consecutive days with tamoxifen and harvested seven days after (all other tissues). Specificity of anti-pERK staining was confirmed by staining tissue sections from EGF-treated wild-type mice (positive control, third row), tissue sections from EGF-treated wild-type mice incubated with phosphatase prior to staining (negative control, fourth row), and tissue sections from wild-type mice treated with the MEK inhibitor PD0325901 (negative control, fifth row). Scale bars = 25 μm.

**Figure S7.**
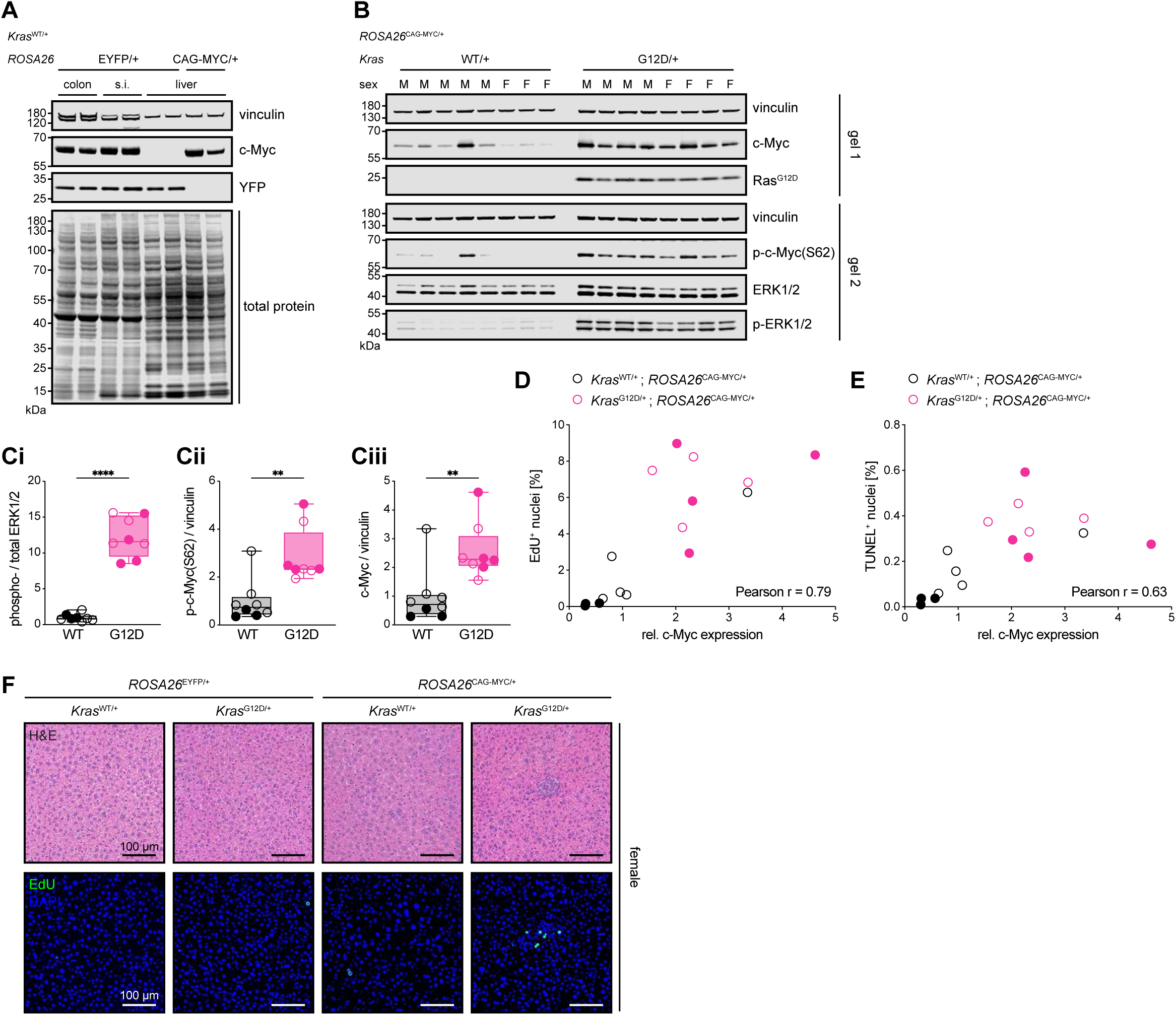
Ectopic expression of c-Myc creates a K-Ras permissive environment in the liver, related to Figure 6. (A) C-Myc protein expression levels in the colon and small intestine of UBC-CreER;R26-YFP;KrasWT mice treated with six doses of tamoxifen and harvested after seven days compared to c-Myc expression levels in livers of R26-YFP;KrasWT and R26-MYC;KrasWT mice treated with 2×10^11^ GCs of AAV8.Cre and harvested 13 days after. (B) ERK activation and abundance of total and phosphorylated c-Myc in livers from R26-MYC;KrasWT and R26-MYC;KrasG12D mice treated with 2×10^11^ GCs of AAV8.Cre and harvested 13 days after. (C) Quantification of Western blot shown in (B). Relative abundance of ERK1/2 phosphorylated at Thr202/Tyr204 normalized to total ERK1/2 (Ci). Relative abundance of c-Myc phosphorylated at S62 (Cii) and total c-Myc (Ciii) normalized to vinculin. Box plots indicate 25^th^ to 75^th^ percentile and group median. Statistical significance between group means was determined by Welch’s t-test. **** P < 0.0001, ** P < 0.01. (D) For each individual liver sample relative c-Myc expression as determined in (Cii) is plotted against its proliferative index indicated by the % of EdU^+^ nuclei. (E) Relative c-Myc expression in individual liver samples plotted against the apoptotic index indicated by % of TUNEL^+^ nuclei. (C-E) Samples from male mice are shown as empty circles, samples from female mice as filled circles. (D and E) Pearson correlation coefficients between c-Myc expression and the second variable are depicted at the bottom right. (F) Representative liver images from female mice ectopically expressing either EYFP or c-Myc and K-Ras^WT^ or K-Ras^G12D^ and harvested after 28 days. Proliferating cells are marked by anti-EdU staining. Scale bars = 100 μm.

